# miR-26 deficiency causes alterations in lens transcriptome and results in adult-onset cataract

**DOI:** 10.1101/2024.01.29.577818

**Authors:** Anil Upreti, Thanh V. Hoang, Minghua Li, Jared A. Tangeman, David S. Dierker, Brad D. Wagner, Panagiotis A. Tsonis, Chun Liang, Salil A. Lachke, Michael L. Robinson

## Abstract

**Purpose:** Despite strong evidence demonstrating that normal lens development requires regulation governed by miRNAs, the functional role of specific miRNAs in mammalian lens development remains largely unexplored.

**Methods:** A comprehensive analysis of miRNA transcripts in the newborn mouse lens, exploring both differential expression between lens epithelial cells and lens fiber cells and overall miRNA abundance was conducted by miRNA-seq. Mouse lenses lacking each of three abundantly expressed lens miRNAs: miR-184, miR-26 and miR-1 were analyzed to explore the role of these miRNAs in lens development.

**Results:** Mice lacking all three copies of *miR-26* (*miR-26^TKO^*) developed postnatal cataracts as early as 4-6 weeks of age. RNA-seq analysis of neonatal lenses from *miR-26^TKO^* mice exhibited abnormal reduced expression of a cohort of genes found to be lens-enriched and linked to cataract (*e.g. Foxe3*, *Hsf4*, *Mip*, *Tdrd7,* and numerous crystallin genes), and abnormal elevated expression of genes related to neural development (*Lhx3, Neurod4, Shisa7, Elavl3*), inflammation (*Ccr1, Tnfrsf12a, Csf2ra)*, the complement pathway, and epithelial to mesenchymal transition (*Tnfrsf1a, Ccl7, Stat3, Cntfr*).

**Conclusion:** miR-1, miR-184 and miR-26 are each dispensable for normal embryonic lens development. However, loss of miR-26 causes lens transcriptome changes and drives cataract formation.

## Introduction

Gene regulation occurs at many levels. Within the lens, many studies have documented the importance of particular transcription factors for establishing lens cell fate and for driving the expression of crystallins, the major proteins expressed by differentiated fiber cells^1^. However, post-transcriptional regulation of gene expression, though less studied in the context of lens biology, also has an important role in lens development and homeostasis^2^. MicroRNAs (miRNAs) play a well-recognized role in post-transcriptional gene regulation, but the role of miRNAs in lens development is not well understood^3,4^.

Several reports suggest that miRNAs may play a role in lens development and pathogenesis^5^. The mouse lens expresses the transcript for miRNA processing enzyme DICER^5,6^, and several miRNA are highly enriched and exhibit spatial and temporal specificity in the lens, raising the question of how these miRNAs function in lens development^7–9^. Targeted deletion of *Dicer* in the mouse lens globally inhibited miRNA processing and led to severe lens degeneration characterized by increased apoptosis and decreased cell proliferation subsequent to E12.5^3,10^. The mouse lens epithelium expresses *miR-204*^9^, and global knockdown of miR-204 in medaka fish resulted in microphthalmia and abnormal lens formation^11^. In the lens, PAX6-induced the expression of *miR-204* resulting in the down-regulation of *Sox11*, *Myo10* and *Fbn2*^4^. Of these three *miR-204* targets, *Sox11*^12,13^ and *Fbn2*^14^ play a role in normal lens morphogenesis. Posterior Capsular Opacification (PCO), the major complication of human cataract surgery, results from an epithelial to mesenchymal transition (EMT) of lens epithelial cells that remain following the removal of the cataractous lens. *miR-204* can inhibit lens cell EMT by negatively regulating SMAD4 in the TGF-β signaling pathway^15^. Notably, *miR-204* expression decreases in PCO^15,16^. Further, deficiency of a cataract-linked gene *Tdrd7* results in misexpression of a cohort of miRNAs that are associated with mRNA targets relevant to lens biology and pathology^17^. In addition, the expression of many different miRNAs changes during fiber cell differentiation^10^ or during cataract development^18–21^.

Despite several studies documenting the expression of miRNAs in the whole lens^7,9,17,20,22^ and lens epithelial cell lines^23^, a clear picture of differential expression of miRNAs in isolated lens epithelium versus lens fiber cells is lacking. To address this gap in knowledge, we conducted a small RNA-seq analysis of newborn mouse lens epithelium and lens fiber cells to quantify and characterize the differential expression of the miRNAs therein. This analysis identified miR-184, miR-26, miR-204, and miR-1 as the most abundantly expressed miRNAs in the newborn mouse lens. To functionally assess their role, we examined lens development in the absence of miR-184, miR-26 or miR-1 in mice. Although mice lacking any one of these miRNAs exhibited normal embryonic lens development, mice lacking miR-26 consistently exhibited aberrant lens gene expression and developed postnatal cataracts.

## Methods

### Animals

All procedures were approved by the Miami University Institutional Animal Care and Use Committee and complied with the ARVO Statement for the Use of Animals in Research, consistent with those published by the Institute for Laboratory Animal Research (Guide for the Care and Use of Laboratory Animals). *FVB/N* mice were euthanized by CO_2_ asphyxiation followed by cervical dislocation. Single and double *miR-1-1* and *miR-1-2* newborn knockout samples were kindly provided by Dr. Deepak Srivastava from University of California, San Francisco.

### Small RNA sequencing, mRNA sequencing and library preparation

Newborn *FVB/N* strain mouse lenses were dissected into capsules containing adhering epithelial cells and fiber cells. Epithelial and fiber cell fractions were each pooled into three biological replicates for a total of six samples, each containing tissue from 8 lenses. Total RNA was extracted using mirVana^TM^ miRNA isolation kit (# AM1560, ThermoFisher Scientific). Small RNA was isolated from total RNA by size-selection and NEBNext Multiplex Small RNA Library Prep Kit was used for 50bp-single-ended sequencing to yield ∼5 million reads per sample.

Four whole lenses were harvested from miR26 triple knockout (TKO) mice at two stages (Day 5 and 20 weeks). For 20-week-old mice we selected lens exhibiting cataract (C) and TKO mice without cataract upon visual inspection. RNA extraction was performed using RNEasy Mini Kit (Qiagen cat. 74104) following the manufacturer’s instructions. The RNA Integrity Number (RIN) was determined using an Agilent 2100 Bioanalyzer and samples with RIN > 7 were used for sequencing. RNA passing in-house quality control were sent to Novogene (Sacramento, CA, USA) for mRNA library preparation and sequencing using NovaSeq PE150 with approximately 30 million reads per sample.

### RNA Seq Data Analysis

Raw reads were quality-analyzed using FastQC (Babraham Bioinformatics-FastQC A Quality Control tool for High Throughput Sequence Data) and MultiQC. Low-quality bases and adapters were trimmed using Cutadapt 3.4 and Trim Galore 0.6.5 with the parameters -q 20 --phred33 --length 20. Mouse genome GRCm39 version: M27 was indexed using Hisat2 (2.1.0-4), incorporating splice junctions from Gencode GTF gencode.vM27.annotation.gtf file^24^. Gene counts were generated using Stringtie2.1.5^25^ and gencode GTF annotation gencode.vM27.annotation.gtf. Differential expression testing was performed with DESeq2^26^. Differentially expressed genes (DEGs) are defined throughout by an adj. *p*-value < 0.05, and log_2_ fold change (LFC) ≥1 criteria were applied.

For microRNA-sequencing, bowtie (version1.3.1) was used for alignment followed by mirdeep2 (0.1.2)^27^ and the miRbase release 22.1. Gene ontology analysis was done using gPRofiler^28^. Pathway enrichment analysis was performed with the Database for Annotation, Visualization and Integrated Discovery (DAVID) online tool^29,30^. Venn diagrams were made using Venny^31^.

### miRNA target prediction

miRNA target prediction was done using two software miRwalk^32^ and Targetscan (8.0)^33^. Only the targets predicted by both of the software were used for downstream analysis. All default parameters were used.

### Gene set enrichment analysis (GSEA)

Gene set enrichment analysis (GSEA) was performed on the normalized count matrix obtained from DESeq2, and murine genes were converted to human orthologs. GSEA was performed using 1000 permutations and gene set permutations with gene set size filters; min = 15 and max = 500. Hallmark gene set was used for analysis^34^.

### Quantification by RT-qPCR

For mRNA genes, cDNA was synthesized using Superscript III Reverse Transciptase (#18080044, ThermoFisher Scientific. qPCR assays were performed on the cDNA using Gotaq Green Master Mix (Promega) following the manufacturer’s instruction and read using CFX96 connect (BioRad). Intron-spanning primers were designed to specifically quantify targeted mRNA transcripts. Each biological sample was analyzed in triplicate by qPCR. The cycling conditions consisted of 1 cycle at 95° C for 100s for denaturation, followed by 40 three-step cycles for amplification (each cycle consisted of 95° C incubation for 20s, an appropriate annealing temperature for 10s, and product elongation at 70° C incubation for 20s). The melting curve cycle was generated after PCR amplification. The reaction specificity was monitored by determination of the product melting temperature, and by checking for the presence of a single DNA band on agarose gels from the RT-qPCR products. Gene expression was calculated and normalized to GAPDH level using delta-delta Ct method (Applied Biosystems). Full primer sequences were listed on Table S4. The expression level of miRNAs was quantified using specific Taqman probes (ThermoFisher scientific) following the manufacturer’s instruction and normalized to snoRNA-202 level. Statistical analysis of RT-qPCR data was performed using student two tailed T test on 3 or more independent experiments. Error bars represent SEM. Differences were considered significant when *p value ≤ 0.05.

### Generation of miR KO mice

Knockout (KO) mice for *miR-184* and for each member of the *miR-26* family were generated using CRISPR/Cas9 via zygote microinjection. Two specific gRNAs flanking each miRNA genomic sequence were designed using CRISPR design tool (http://crispr.mit.edu/) and synthesized via gBlocks Gene Fragment (IDT integrated DNA Technologies). gRNAs were *in vitro* transcribed using *in vitro* MEGAscript T7 transcription kit (#AM1334, ThermoFisher Scientific) and purified using MEGAclear Transcription Clean-up kit (#AM1908, ThermoFisher Scientific). A mixture of Cas9 mRNA (50 ng/ µl, #L-6125, TriLink Biotechnologies) and two specific gRNAs (25 ng/ µl each) for each miRNA target were injected to single-cell zygotes. Desired miR KO mice were screened by PCR and confirmed by DNA sequencing and RT-qPCR (primers listed in Table S3. Compound *miR-26* KO mice were generated by inter-crossing single KO mice for each of *miR-26a1, miR-26a2* and *miR-26b*. To test the off-targeted effects of the gRNAs, for each gRNA, the top 4 genes with highest risk of being targeted in the exon regions in founder mice were analyzed and confirmed by PCR and DNA sequencing.

### Lens photography

Age-matched animals were euthanized and their eyes were dissected in phosphate-buffer saline (PBS). Lenses were dissected in PBS and photographed using a Motic Stereo Zoom microscope.

### Histology

Tissues were collected and fixed in 10% neutral buffered formalin for 24 hr. Standard protocols were used to process and embed tissues in paraffin wax before sectioning at 5μm thickness. Standard hematoxylin and eosin-stained sections were performed to analyze the structure of the lens, and images were captured using a Nikon TI-80 microscope.

## Results

### small RNA-seq facilitates comparative expression of miRNAs between lens epithelial and fiber cells

We collected RNA from isolated lens epithelial and fiber cells from newborn mice and performed small RNA-seq to analyze the differential expression of miRNAs. A distance matrix of expressed miRNAs clearly distinguished lens epithelial cells from lens fiber cells (Figure S1). Of all annotated miRNAs that were detected in the lens, 184 were differentially expressed between epithelial and fiber cells (fold change >1, adjusted p value <0.05) (Figure 1A). Lens epithelial cells were enriched for 76 miRNAs and fiber cells were enriched for 108 miRNAs (Table S1). The top 25 differentially expressed miRNAs (Figure 1B), included 16 and 9 miRNAs that were more abundantly expressed in fiber cells and epithelial cells, respectively.

**Figure 1.**
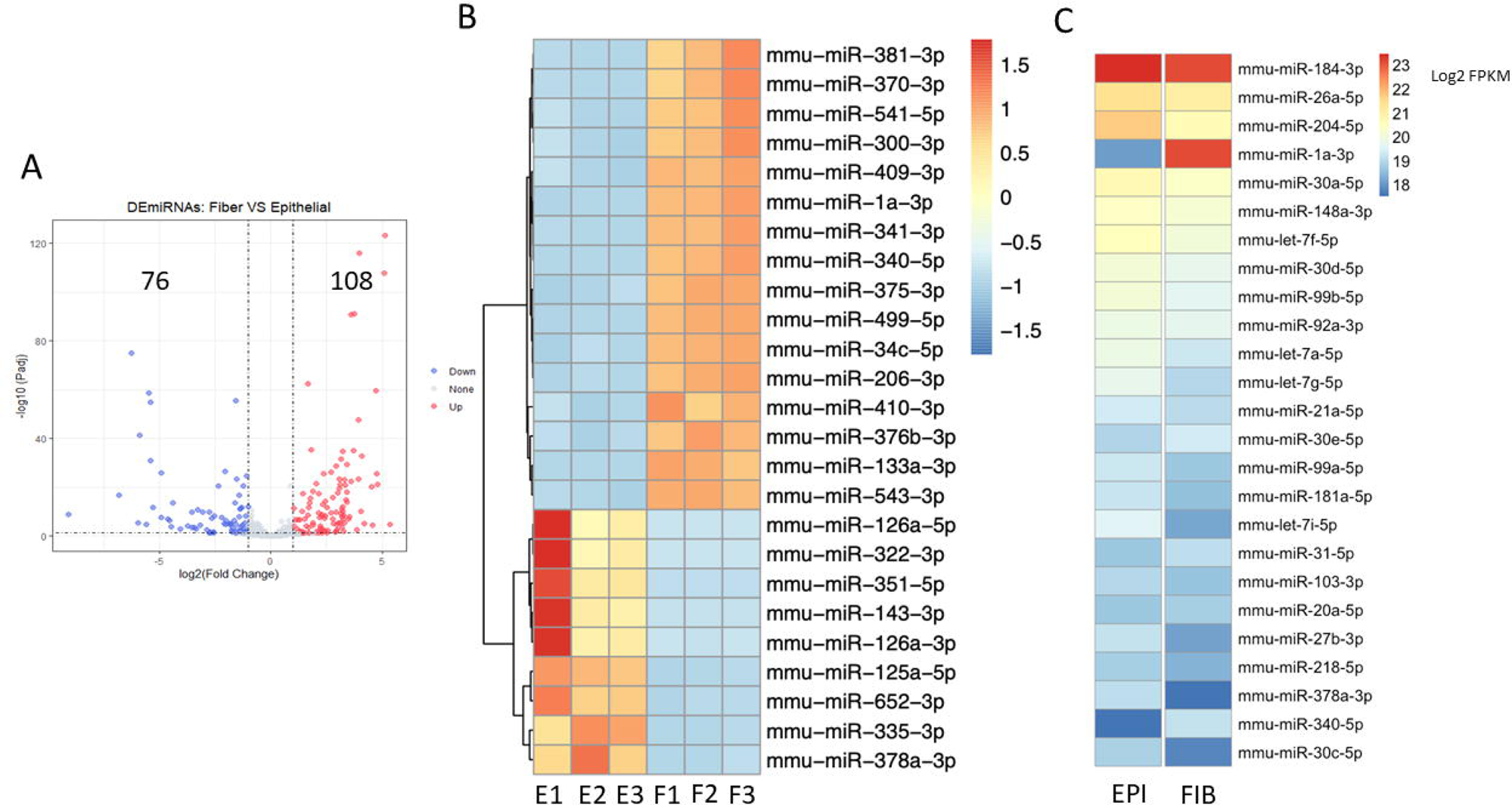
miRNA profiling of newborn lenses. (A) Volcano plot representing the differentially expressed genes between lens epithelium and lens fiber cells. Red represents the up-regulated genes (enriched in fiber cells) and blue represents the down-regulated genes (enriched in epithelial cells). (B) Z-score heatmap displays top 25 differentially expressed miRNAs in lens epithelium and lens fiber cells. E: epithelial cells, F: fiber cells (C) Comparative average expression of the most highly expressed miRNAs in lens epithelium and fiber cells. Transcript level was expressed as miRNA counts normalized transformed by log_2_(FPKM) and plotted in a heatmap.

The lens-expressed miRNAs were also examined in terms of overall abundance across both epithelial and fiber cells by averaging the log_2_ fragments per kilobase of transcripts per million mapped reads (FPKM) from all six samples. The top 25 miRNAs, listed in terms of overall abundance in the lens, are shown in a heatmap based on their expression in lens fiber cells (Figure 1C). Three of the top differentially expressed miRNAs (miR-1, miR-340, and miR-378a) were also found in the 25 most abundantly expressed miRNAs. The most abundant miRNAs in the lens included miR-184, miR-26a, miR-204, and miR-1 (Table S2). Focusing on these most abundant miRNAs in the lens, miR-184 exhibited high expression in both cell types. miR-26a transcripts were not differentially expressed between the epithelium and fibers. miR-204 was expressed significantly higher in the lens epithelium than in the lens fibers. In contrast, fiber cells expressed significantly more miR-1 than epithelial cells. Given that multiple lines of *miR-204* knockout mice have been reported (see discussion), we chose to focus on miR-184, miR-1 and miR-26. The expression of these microRNAs in lens epithelial cells and lens fiber cells was confirmed by RT-qPCR (Figure S2). Given their relatively high expression in the lens, we undertook a functional analysis of miR-184, miR-1 and miR-26 during mouse lens development.

### The role of miR-26 in the lens

To abrogate miR-26 transcripts from the mouse lens, we employed a CRISPR/Cas9 editing strategy. *miR-26* is present in three copies in both the mouse and human genome. The three members of miR-26 gene family include *miR-26a1*, *miR-26a2* and *miR-26b*, each of which are found within introns of three different *Ctdsp* host genes located on three different chromosomes (Figure 2A). While both *miR-26a1* and *miR-26a2* genes produce an identical mature miR-26a-5p, the *miR-26b* gene expresses mature miR-26b-5p. Mature miR-26a-5p and miR-26b-5p sequences share an identical seed region and only differ in two nucleotides, suggesting that they are likely to function redundantly. Each *miR-26* locus was individually targeted by two gRNAs (Table S3) via zygote microinjection (Figure S3). PCR screening showed targeted deletion for each of *miR-26* family members (Figure 2B). DNA sequencing confirmed the loss of almost the entire *miR-26a1* and *miR-26a2* genomic sequences (Figure 2C). However, a much smaller deletion in *miR-26b* was achieved that only disrupted the seed sequence. Among the four highest scoring potential off-targeted genes for each gRNA, our analysis only showed a small deletion in one allele of *Tbk1* by the miR-26a1_gRNA2, which was eliminated by backcrossing to *FVB/N* wild-type mice.

**Figure 2.**
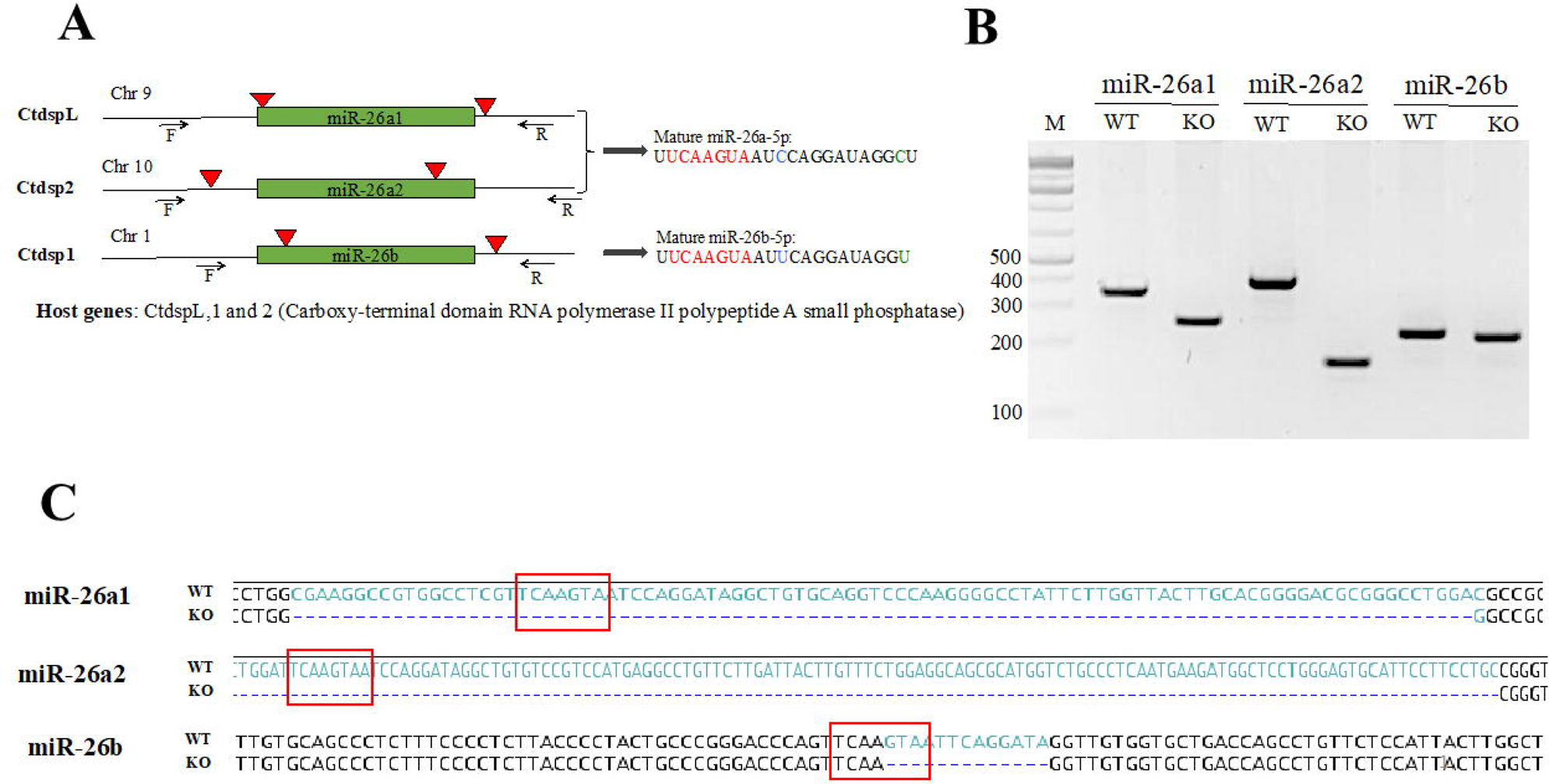
Generation of mice deficient in members of miR-26 family. (A) A diagram illustrates the generation of miR-26 family KO mice using CRISPR/Cas9 technology. Three members of the miR-26 family are located within the *Ctdsp* host gene family, and produce mature miRNAs with identical seed sequences (red highlight). A mixture of Cas9 mRNA and 2 gRNAs specific for each miRNA gene were injected into zygotes via microinjection. Arrowheads represent the cutting sites of Cas9 enzymes. Arrows represent the locations of the forward (F) and reverse (R) PCR primers used for genotyping of resultant mice. Chr = Chromosome. (B) PCR with specific primers was used for screening for targeted deletions in tail DNA from homozygous knockout (KO) mice, resulting in expected reductions in amplicon size, relative to that in DNA from wild-type (WT) mice. (**C**) DNA sequencing analysis demonstrated the deletions in the miR-26 sequences with complete loss of *miR-26a1* and *miR-26a2* seed sequences and the partial loss of the *miR-26b* seed sequence (red boxes) in the respective KO mice. The deleted nucleotides in the KO mice are represented by green text in the wild-type (WT) sequence.

Each targeted deletion of *miR-26a1* or *miR-26a2* led to a significant reduction of mature miR-26a-5p production in single knock-out (KO) lenses (Figure 3A-B). The small deletion in *miR-26b* eliminated miR-26b-5p production (Figure 3C). To determine whether targeted disruption of *miR-26a1, miR-26a2* and *miR-26b* affected the expression of their host genes, we performed RT-qPCR on lens RNA for transcripts for *CtdspL, Ctdsp1* and *Ctdsp2* in the context of the relevant *miR-26* KO (Figure S4). In no case, did the deletion of a copy of miR-26 significantly affect the expression of the *Ctdsp* host gene.

**Figure 3.**
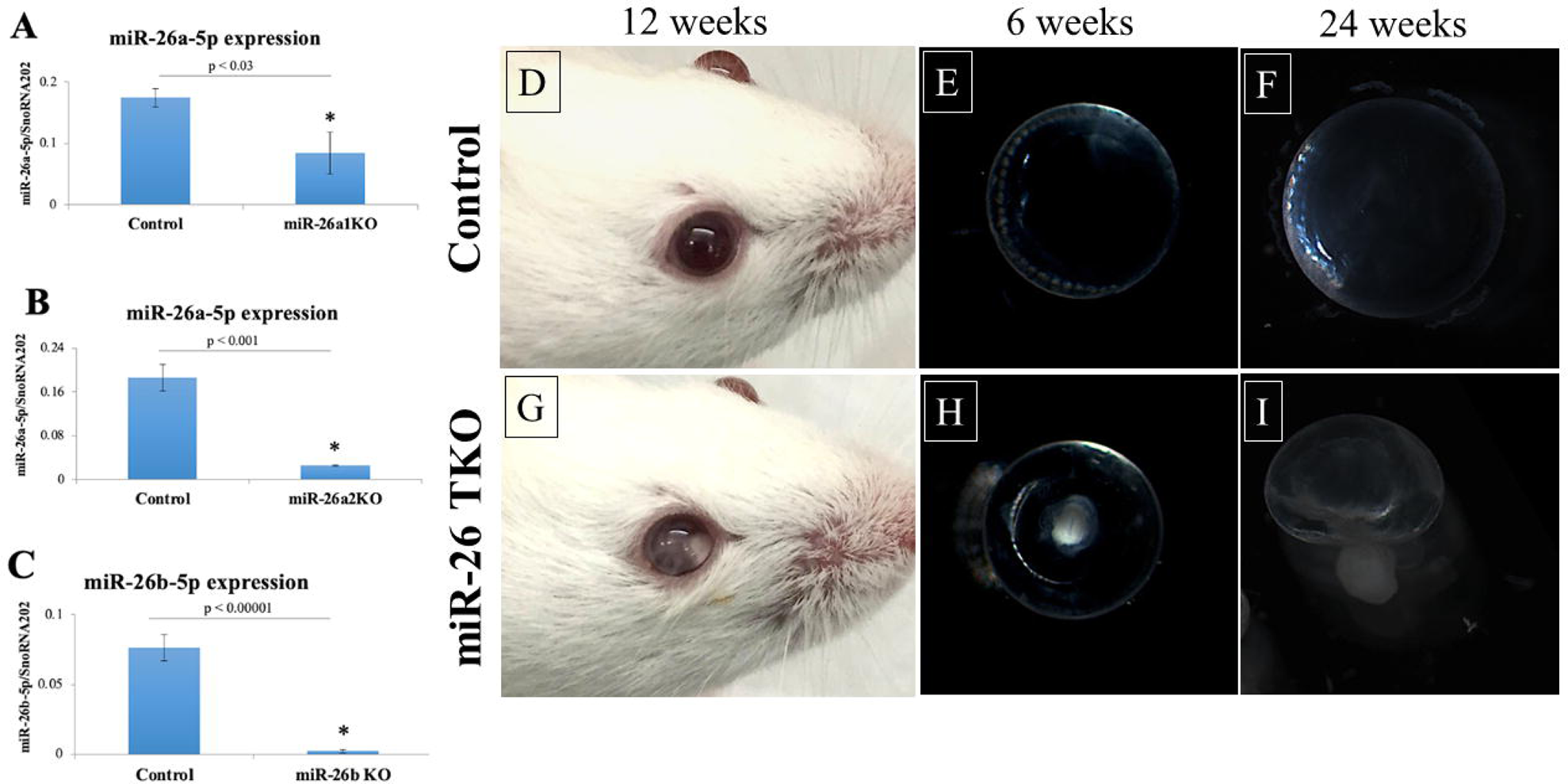
Severe cataract in adult *miR-26^TKO^* mice. (A-B) A significant reduction of mature miR-26a-5p level in 3-week-old single *miR-26a1* and *miR-26a2* KO lenses was observed, assessed via RT-qPCR. (C) miR-26b-5p expression was completely abolished in 3-week-old miR-26b KO lenses. (D,G) Cataract was observed in *miR-26^TKO^*mice at 12 weeks old as compared with the control mice. (E,H) *miR-26^TKO^*lenses showed smaller size and apparent nuclear cataract at 6 weeks old as compared with the control. (F,I) At 24 weeks old, TKO lenses ruptured and were severely deformed. Error bars on the graph represent SEM and the asterisk represents a significant difference from the control value. N.S, no significance.

Mice homozygous for any of the single *miR-26* KO genes (*miR-26a1, miR-26a2,* or *miR-26b*) were viable, fertile and without any obvious lens phenotype. Likewise, lenses from any combination of double *miR-26* KO alleles appeared normal (data not shown). However, mice homozygous for all three miR-26 deletions (*miR-26a1, miR-26a2,* and *miR-26b*), hereafter referred to as *miR-26^TKO^* mice, developed nuclear cataracts as early as 4 weeks of age, with 75% (9/12) of *miR-26^TKO^* mice displaying cataracts in at least one eye by 6 weeks of age (Figure 3D-I). Often, the cataracts started unilaterally, but eventually both eyes developed cataracts such that by 22 weeks of age, bilateral cataracts had developed in 100% of the mice (N=10). These cataracts progressed with time such that by 24 weeks, most lenses had ruptured through the capsule (Figure 3I). Although all of the single *miR-26* KO mice exhibited normal fertility, increasing the number of *miR-26* KO alleles had a negative effect on fertility. *miR-26^TKO^* mice typically become infertile after one or two litters, with reproductive tract tumors often appearing in *miR-26^TKO^*males (data not shown). As a result, *miR-26^TKO^* mice were preferably generated by mating mice homozygous for deletions in two *miR-26* alleles and heterozygous for the third *miR-26* allele.

### Transcriptome changes in miR-26 TKO mouse lenses

To gain mechanistic insight into lens pathology in *miR-26^TKO^*mice, we performed RNA-seq on *miR-26^TKO^* lenses at two stages: five days after birth (P5), well before the appearance of cataracts, and at twenty weeks (W20), a time at which most *miR-26^TKO^* mice had developed cataracts. For the W20 stage, we collected RNA from *miR-26^TKO^*lenses with cataract (C) and without an obvious cataract (NC). We compared gene expression in these *miR-26^TKO^* samples to the gene expression in age-matched wild-type control (*FVB*) lenses. Distance matrix clustering (pairwise comparisons of total gene expression from each sample) revealed distinct clustering of replicates within each experimental group (Figure 4A). The *miR-26^TKO^* and *FVB* lenses at P5 showed the closest global relationship among the analyzed groups via hierarchical clustering; however, the W20 *miR-26^TKO^* lenses were closer in overall gene expression to W20 *FVB/*wildtype lenses than the P5 *FVB* lenses. A 3-dimensional principal component analysis plot (Figure 4B) demonstrated close clustering of replicates from each experimental group. Consistent with the distance matrix, the P5 *FVB* and *miR-26^TKO^*samples displayed a relatively clustered spatial proximity as opposed to the W20 samples. Similarly, W20 *miR-26^TKO^* lenses with cataract formed a cluster notably distinct from all other groups. Interestingly, these data underscore that the transcript profile of lenses presenting both NC and C are quantifiably distinct, even when collected at the same age and if collected from contralateral eyes of the same mouse.

**Figure 4.**
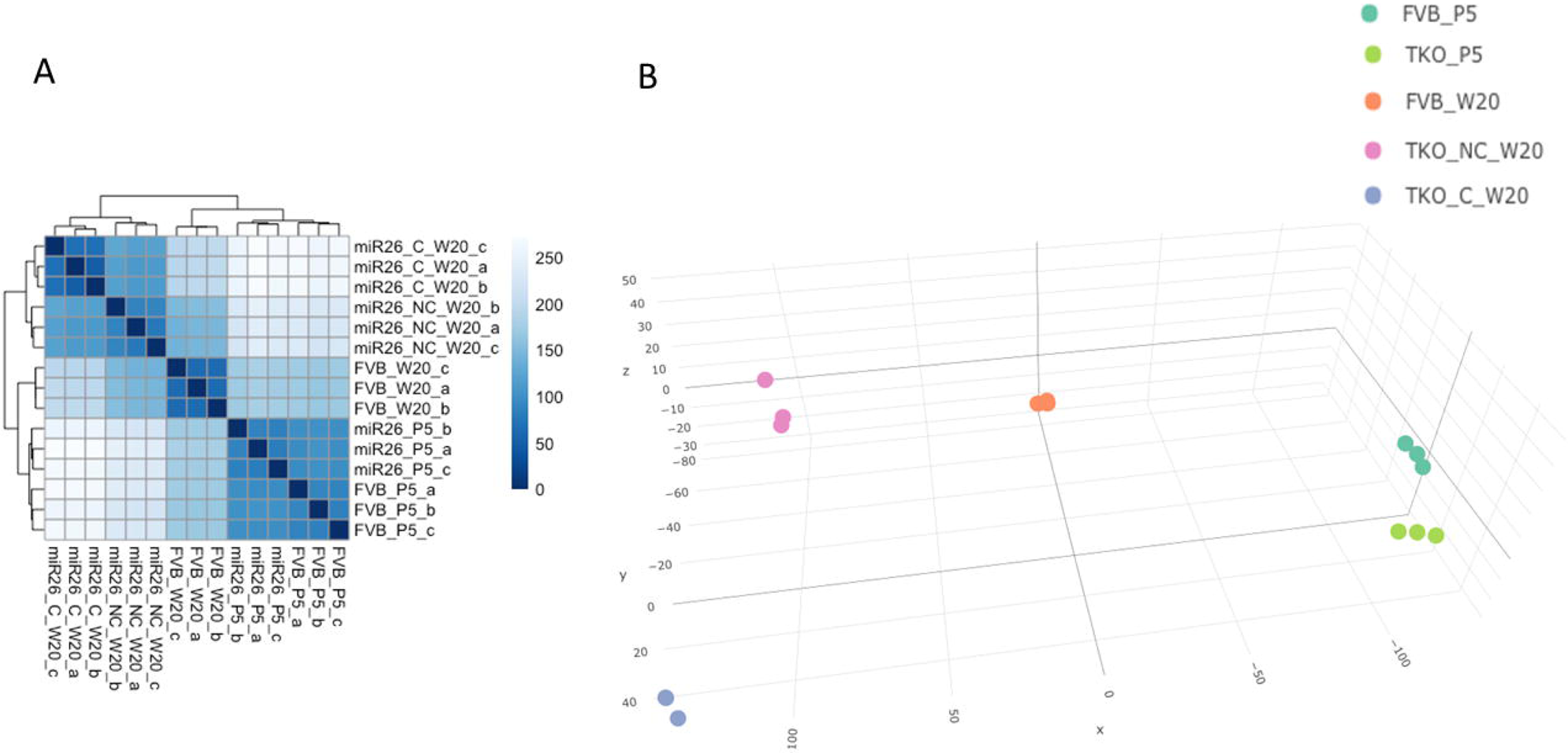
mRNA profiling of *miR-26^TKO^* mice at different stages. (A) Distance matrix indicates difference between wild type (*FVB/N*) and miR-26 knockout at five day (P5), twenty week old mice with cataract (TKO_C_W20) and twenty week old mice without cataract (TKO_NC_W20). (B) A three-dimensional principal component analysis plot shows tight clustering of the three replicates within each group.

A total of 1,653 genes (1,000 up-regulated and 653 down-regulated) exhibited differential expression (log_2_ fold change ≥1, p_adjust_ <0.05) between the normal *FVB* and *miR-26^TKO^* lenses at P5, before the onset of cataract development. With age, the number of differentially expressed genes increased between *FVB* and *miR-26^TKO^* lenses. At W20, 5,143 genes (1,476 up-regulated and 3,667 down-regulated) were differentially expressed between the *FVB* lenses and the *miR-26^TKO^* lenses that did not exhibit overt lens opacity at this age. This differential gene expression number at W20 rose to 8,241 (3,171 up-regulated and 5,070 down-regulated) when comparing the *miR-26^TKO^* lenses with cataracts to the *FVB* lenses. All relevant differential gene comparisons are listed in Table 1.

**Table 1:** Differential Gene Expression Analyses Between Wild-Type (FVB) and miR-26 Triple Knockout (TKO) Lenses.

| Conditions | DEGs | Up | Down | Table Number |
| --- | --- | --- | --- | --- |
| FVB_P5_vs_TKO_P5 | 1653 | 1000 | 653 | S5 |
| FVB_W20_vs_TKO_C_W20 | 8241 | 3171 | 5070 | S6 |
| FVB_W20_vs_TKO_NC_W20 | 5143 | 1476 | 3667 | S7 |
| TKO_P5_vs_TKO_NC_W20 | 8851 | 2529 | 6322 | S8 |
| TKO_P5_vs_TKO_C_W20 | 10287 | 3453 | 6834 | S9 |
| FVB_P5_vs_FVB_W20 | 6264 | 2040 | 4224 | S10 |

W20 = Week 20, P5 = Postnatal day 5, C = Cataract, NC = No Cataract, DEGs = Differentially Expressed Genes

To determine the effect of miR-26 loss on lens fiber cell differentiation, we compared genes typically associated with lens epithelial cells (Figure 5A) or lens fiber cells (Figure 5B) in each condition. The pattern of epithelial gene expression segregated into six main groups (I-VI). Genes in group I exhibited relatively low expression in both *FVB* and *miR-26^TKO^* lenses at P5 and in *FVB* lenses at W20. However, these genes exhibited abnormally elevated expression in the *miR-26^TKO^* lenses at W20, with the highest expression seen in lenses with obvious cataracts. Genes in group I include two VEGF receptor genes, *Flt1* and *Kdr*, *Gabbr1* (encoding the GABA B1 receptor), *Dach2* (a transcription factor), and *Cx3cl1* (a chemokine associated with neurons and glia). The expression pattern of group II genes exhibited reduced expression in the P5 *miR-26^TKO^* lenses relative to the FVB lenses. All of the group II genes were more highly expressed at W20 with *Dll1* showing peak expression in *FVB* lenses, *Pdpn* showing peak expression in *miR-26^TKO^* lenses without cataract and *Rgs6* showing peak expression in *miR-26^TKO^*lenses with cataract. Genes in group VI (*Slc22a23, Slc38a3* and *Sulf1*) were expressed at an intermediate level in both FVB and *miR-26^TKO^*lenses at P5, exhibited very low expression in *FVB* lenses at W20, and reached their highest expression in W20 *miR-26^TKO^* lenses with cataracts. The genes in group III (including *Npnt, Cdh1, Foxe3* and *Pdgfra*) were expressed at the highest level in both P5 samples with most of these genes showing reduced expression in the *FVB* lenses at W20. Importantly, these key genes showed reduced expression in W20 *miR-26^TKO^* lenses without cataracts and even further reduction in expression in the W20 *miR-26^TKO^*lenses with cataracts. Genes in group IV exhibited generally higher expression in *FVB* lenses at W20 with mild and marked reductions in expression in W20 *miR-26^TKO^*lenses without and with cataracts, respectively. The genes in group V (including *Cdk1, Mki67* and *Btg1* associated with cell proliferation) exhibited peak expression in the P5 samples with generalized reductions in expression at WK 20. In sum, key lens epithelial genes (*e.g.*, *Cdh1*, *Foxe3*, *etc.*) showed reduced expression in W20 *miR-26^TKO^*lenses with cataract, suggesting that alteration of normal epithelial transcriptome could contribute to the lens defects in these mice.

**Figure 5.**
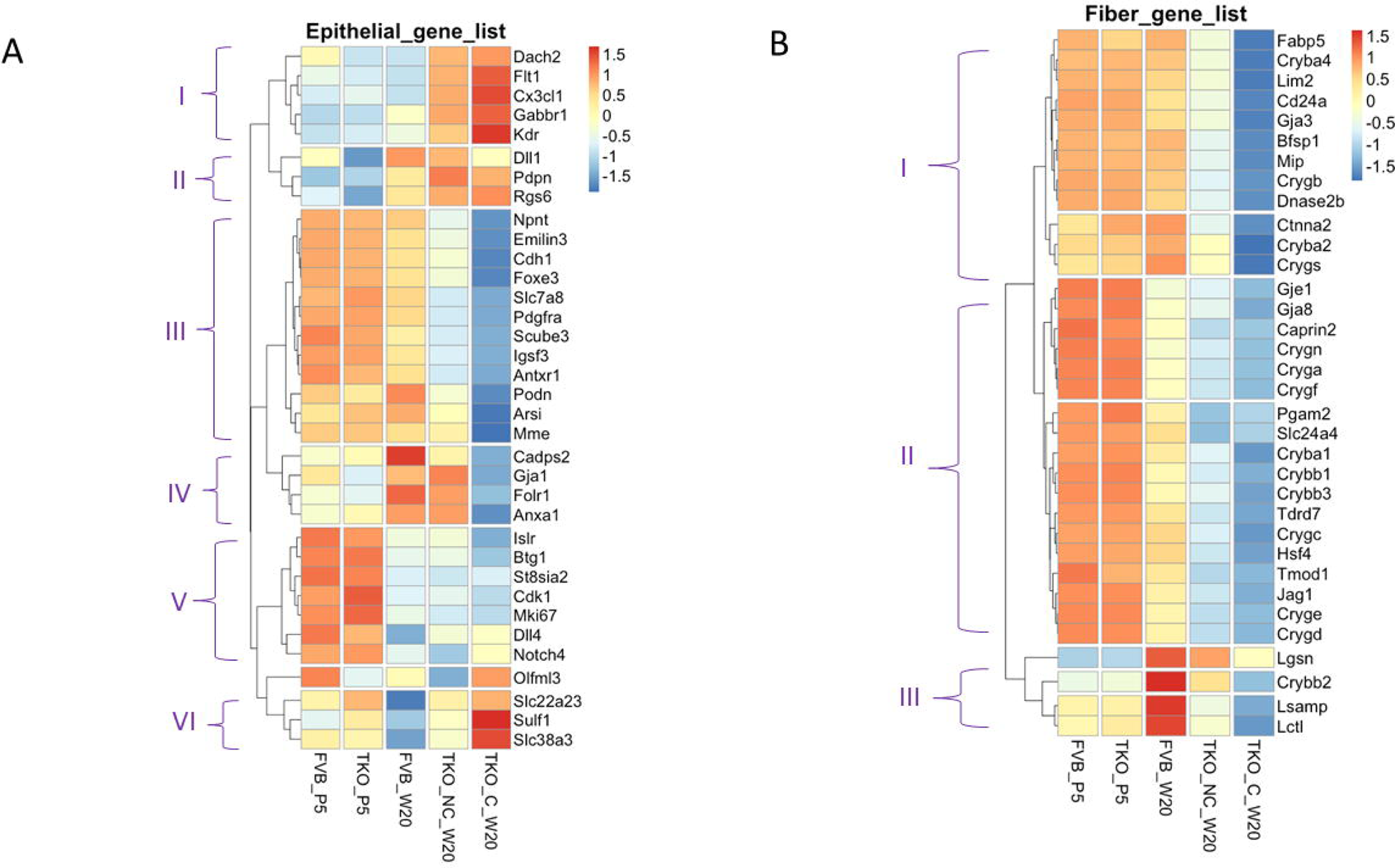
Lens fiber cell differentiation is severely affected in *miR-26^TKO^* mice at later stages (W20). (A) Heatmap indicates z-score adjusted expression values to reveal a clear transition of epithelial genes across tested conditions, indicating the important role of miR-26 in maintaining epithelial cell identity. (B) Heatmap indicates z-score adjusted expression values to show a clear transition of fiber genes across tested conditions, indicating the important role of miR-26 in facilitating fiber cell differentiation.

The pattern of fiber cell gene expression (Figure 5B) fell into three major groups (I-III). The most obvious characteristic shared by all three groups was low expression of several key fiber cell genes in the W20 *miR-26^TKO^* samples, whether with or without cataract. The genes in group I (including *Cryba4, Cryba2, Crygs, Lim1, Gja3, Bfsp1, Mip,* and *Dnase2b*) exhibited high expression in both P5 lens samples (with no change between control and *miR-26^TKO^*) and in W20 *FVB* lenses. However, a significant drop in the expression of these key lens genes was observed in *miR-26^TKO^*W20 samples without cataract, and these were even further reduced in W20 *miR-26^TKO^*lenses with cataract. Genes in group II (including *Gja8, Hsf4, Tmod1* and several genes encoding β- and γ-crystallins) exhibited high expression in both P5 lens samples with progressively decreasing expression in the *FVB* lenses with age. Again, compared to control, *miR-26^TKO^* samples without and with cataract at W20 showed significant reduction in these genes. Finally, Group III genes (including *Crybb2, Lgsn* and *Lctl*) exhibited low expression at P5 with peak expression in the W20 *FVB* lenses and low and very low expression in the *miR-26^TKO^* lenses without and with cataracts, respectively. Thus, while the majority of these fiber genes had normal expression in control and *miR-26^TKO^* at P5, by WK 20, they were significantly reduced in the *miR-26^TKO^* samples.

We also examined genes that were differentially expressed in any of the *miR-26^TKO^* samples and included in the list of genes recognized as cataract-associated in the Cat-Map database^35^ or the list of genes exhibiting high “lens-eneriched” expression and recognized as high-priority in the iSyTE database^36^. Of the 496 genes listed in Cat-Map, 58 (11.7%) of these are up-regulated (Figure S5) and 145 (29.2%) are down-regulated (Figure S6) in the miR-26 *miR-26^TKO^* lenses. Four genes (*Aipl1, Ndp, Shh* and *Otx2*) are up-regulated and two genes (*Bfsp2* and *Myo7a*) are down-regulated in all *miR-26^TKO^* conditions. Of the 528 genes listed in iSyTE as enriched in the lens, 76 (14.4%) are up-regulated (Figure S7) and 290 (54.9%) are down-regulated (Figure S8). Three genes: *Crym, Rrh,* and *Kcnk1* were upregulated in all of the *miR-26^TKO^* samples and three genes: *Bfsp2, Hspb1,* and *Frem2* were down-regulated in all of the *miR-26^TKO^*samples. While there was overlap in both Cat-Map and iSyTE gene lists, in general more genes in both lists were down-regulated than were up-regulated. Genes characteristic of fiber cell differentiation and lens identity (eg. *Bfsp2, Cryga, Crybb2, Dnase2b, Foxe3, Gja8, Mip,* and *Tdrd7*) tended to be down-regulated in the *miR-26^TKO^*samples. In contrast, most of the up-regulated genes the *miR-26^TKO^* samples were related to cellular signaling (eg. *Bmp3, Porcn, Rgs6, Ndp,* and *Shh*) or transcription factors associated with retinal development (eg. *Nrl*, *Otx2*, and *Vsx2*).

### Identification of direct targets of miR-26 in the lens

To identify potential targets of miR-26, we utilized two web based tools (Targetscan and miRWALK) to evaluate all the protein-coding genes in the mouse genome. This analysis identified a total of 8,727 predicted targets and 1,520 of these targets (17.4%) were predicted by both software packages (Figure S9). We analyzed these 1,520 potential target genes for differential expression in the *miR-26^TKO^* lenses. Of these potential targets, 396 (∼26%) were up-regulated (Figure 6A), while 265 (17.4%) were down-regulated in at least one class of *miR-26^TKO^* samples (P5, W20_C or W20_NC) (Table S11). Only 45 (3%) of these potential target genes were up-regulated in the *miR-26^TKO^* lenses at P5, and 26 (1.7%) of these genes were commonly up-regulated in all miR-26 conditions (P5, C_W20 and NC_W20) analyzed (Figure 6B). An additional 352 potential miR-26 targets are up-regulated in the *miR-26^TKO^* lenses at week 20 (C_W20 and/or NC_W20). The 26 commonly up-regulated genes also demonstrate a progressive increase in expression (FVB_P5 < TKO_P5 < FVB_W20 < TKO_NC_W20 < TKO_C_W20), with the *miR-26^TKO^* lenses with cataracts at week 20 showing the highest expression of these genes. 54% of these 26 genes are associated with nervous system development or synaptic membrane proteins (*Acs16, Csf1r, Elavl2, Elavl3, Grik2, Lhx3, Neto1, Neurod4, Shisa7, Snph, Slc1a2, Shank2, Tfap2c,* and *Unc5d*), based on gene ontology analysis. From a gene regulatory standpoint, three of these 26 genes are transcription factors *(Lhx3, Neurod4* and *Tfap2c*), and three others are RNA-binding proteins (*Celf5, Elavl2*, and *Elavl3*). In summary, the majority of the commonly up-regulated predicted miR-26 targets in the lens normally participate in neuronal development or function.

**Figure 6.**
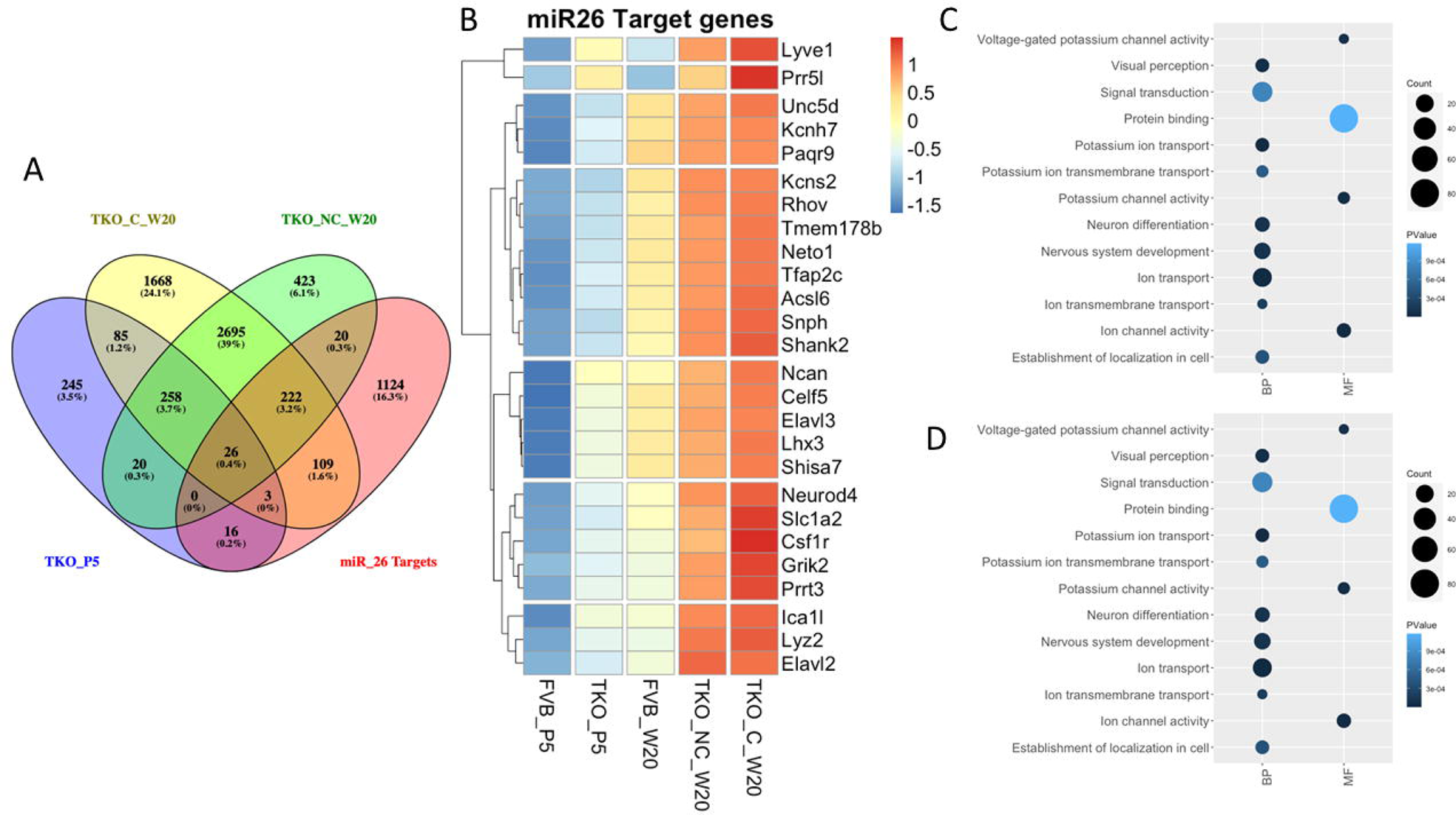
Analysis of predicted miR-26 targets that are up-regulated in *miR-26^TKO^* samples. (A) Venn diagram displays the genes up-regulated in TKO samples intersected with common miR-26 targets predicted by miRwalk and Targetscan. (B) Heatmap displaying z-score adjusted expression values in the lens for the verified twenty-six targets for miR-26 with relative expression in all conditions. (C) Bubble plot represents select gene ontology terms for 258 genes (excluding the 26 genes predicted to be direct miR-26 targets) identified to be differentially expressed in all *miR-26* knockout samples. BP = Biological Processes, MF = Molecular Function. (D) Bubble plot represents top enriched pathways identified using the reactome database for the 258 differentially expressed genes.

There were also 258 genes that were up-regulated in all the *miR-26^TKO^*samples that were not predicted to be direct targets of miR-26. A gene ontology (GO) analysis of these 258 genes (Figure 6C) revealed enrichment for terms relevant to ion transport, neuronal differentiation, and visual perception. When these 258 genes were analyzed for enrichment in the reactome pathway, the top pathways identified included: Neuronal System, Transmission across Chemical Synapses, Potassium Channels and Neurotransmitter release cycle (Figure 6D). Of the 1,520 potential target genes identified, 265 (17.4%) were down-regulated in at least one *miR-26^TKO^* sample (Figure S10 - Table S12). There were 75 genes down-regulated in all *miR-26^TKO^*samples, of which 3 were predicted to be miR-26 targets. Gene ontology analyses of these 75 genes failed to show any significant enrichment in key GO terms. Together, these data suggest that miR-26 normally functions to suppress – in the lens – genes involved in neuronal biology, and thus deficiency of miR-26 may alter the lens transcriptome and contribute to the lens defects. Thus, it appears that most of the up-regulated transcripts in the *miR-26^TKO^* lenses that are not predicted to be direct targets are also primarily involved in neural biology.

### Gene set enrichment analysis of differentially expressed genes in miR-26^TKO^ lenses

Gene set enrichment analysis (GSEA) represents a way to comprehensively explore differential gene expression between any two conditions with respect to molecular signature database collection (hallmark) gene sets which are characteristic of specific biological states or processes. To determine how gene expression changes in the *miR-26^TKO^* mice with age, gene expression in *miR-26^TKO^* lenses at P5 were compared with the *miR-26^TKO^* lenses at W20 without cataract or the *miR-26^TKO^* lenses at W20 with cataract using GSEA. A significant enrichment was observed for genes related to Inflammatory response and Complement in the W20 *miR-26^TKO^*lenses without cataract compared to P5 *miR-26^TKO^* lenses (Figure 7A). Similarly, an enrichment was observed for tumor necrosis factor alpha (TNFA) signaling via NFKB in the *miR-26^TKO^* W20 lenses with cataract compared to the P5 *miR-26^TKO^* lenses. Given these findings, we compared gene expression related to genes listed under Inflammation in the GSEA list in all five conditions. Altogether, there were 63 genes related to inflammation that were differentially expressed (Figure 7B). Most of the inflammation-related genes exhibited low expression in the P5 lenses, moderate to low expression in the *FVB* lenses at W20 and moderately high to high expression in the W20 *miR-26^TKO^* lens samples, with the highest expression in those lenses displaying cataract (including *Ccr1, Tnfrsf12a, Csf2ra,* and *Stat3*). These results underscore broad dysregulation of gene sets involved in inflammation and the complement cascade in *miR-26^TKO^* lenses, which could implicate aberrant immune responses in the observed cataractogenesis.

**Figure 7.**
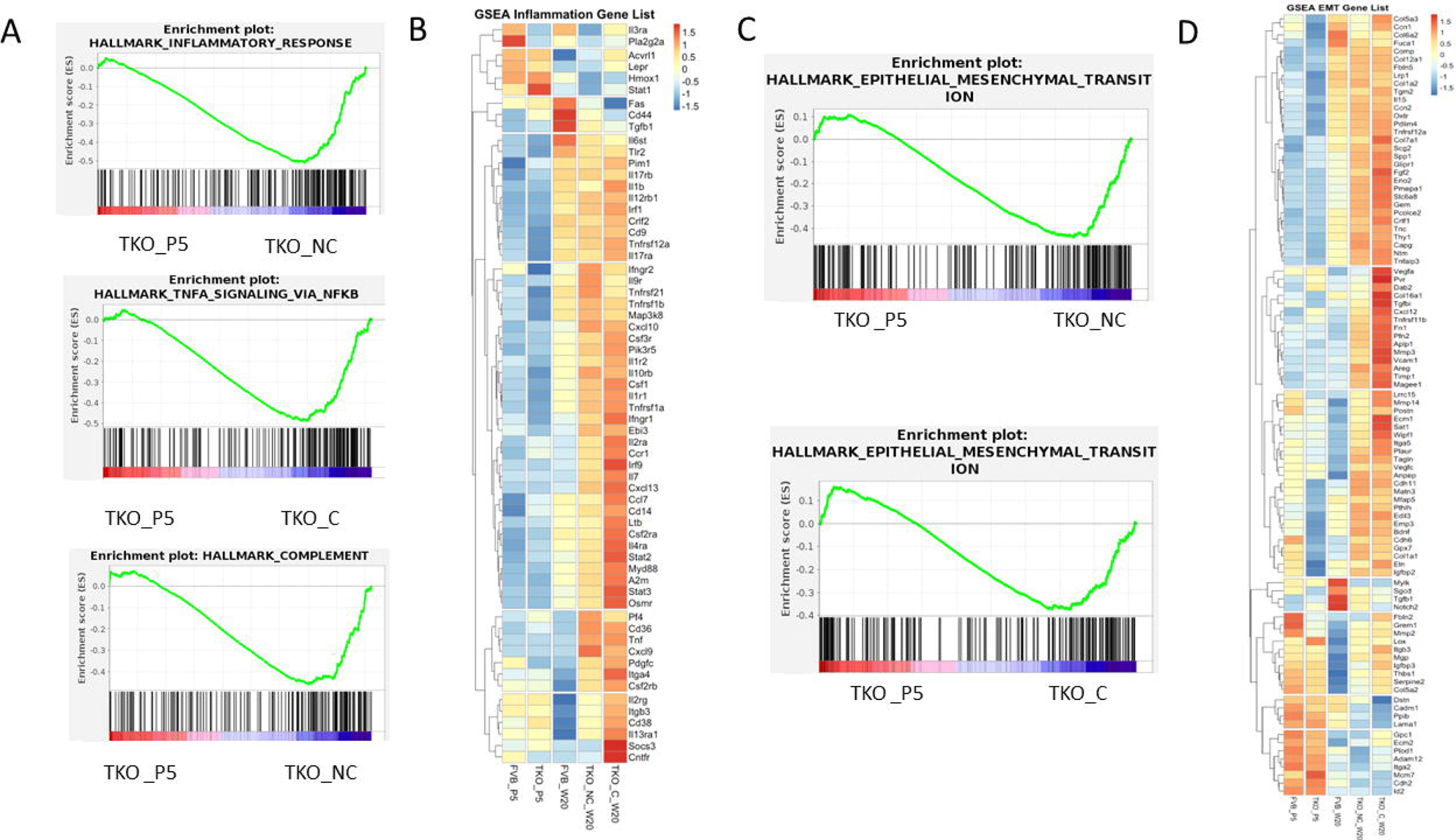
*miR-26* TKO_W20 transcriptomes are enriched for immune response and Epithelial to mesenchymal transition (EMT) fate. (A) GSEA enrichment plot represents the enrichment of the terms inflammatory response, TNFA signaling via NFKB, and complement in TKO_W20_NC and TKO_W20_C samples, respectively when compared with TKO_P5 samples. Genes are ordered along the x-axis based on expression rank between the two conditions. Black bars indicate genes associated with a given term. The green line indicates the enrichment score determined by GSEA. (B) Heatmap using z-score scaled expression values shows inflammation genes across all conditions. (C) GSEA enrichment plot represents the enrichment of Epithelial to mesenchymal transition (EMT) in TKO_W20 samples. (D) Heatmap using z-score scaled expression values shows EMT-associated genes across all conditions.

GSEA analysis also suggested that the *miR-26^TKO^* lenses at week 20 were undergoing significant epithelial to mesenchymal transition (EMT). Increased EMT was a characteristic of both W20 *miR-26^TKO^* lens samples regardless of cataract status compared to the P5 *miR-26^TKO^* lenses by GSEA (Figure 7C). We explored the expression of 95 genes related to EMT that were differentially expressed in our *miR-26^TKO^* lenses. There were significant differences in the expression of these genes between the two P5 samples. In general, the P5 *miR-26^TKO^* lenses exhibited a lower expression of EMT related genes than the P5 *FVB* lenses. In contrast, the majority of these genes (including *Ccn1, Ccn2, Tgfbi, Vegfa*, *Fn, Vcam1, Mmp3,* and *Mmp14*) were expressed most highly in the W20 samples with cataract followed by the W20 samples without cataract and expressed least in the *FVB* W20 samples (Figure 7D). Twenty of the EMT genes (including *Fbln2, Mmp2, Col5a2, Lama1*, and *Cdh2*) were more highly expressed in the P5 *FVB* lenses than in the W20 *miR-26^TKO^*lenses and four of these EMT-related genes (*Mylk, Sgcd, Tgfb1,* and *Notch2*) were most highly expressed in the *FVB* W20 sample. Thus, miR-26 TKO led to temporally-controlled dysregulation of EMT gene sets, lending direct insights to regulators of EMT that may participate in cataract formation.

To gain more insight into early changes in the lenses lacking miR-26, we performed GSEA analysis on differentially expressed genes in the *FVB* and *miR-26^TKO^* samples at P5. This revealed a specific enrichment for genes associated with the G2M checkpoint and E2F targets in the *miR-26^TKO^*lenses (Figure S11A-B). Both of these hallmark gene sets suggest an alteration of cell cycle control in the P5 *miR-26^TKO^* lenses. To further explore cell cycle regulation in the *miR-26^TKO^* lenses, we compared gene expression for E2F target genes in all five conditions (Figure S11C). Almost all of these genes demonstrated peak expression in the *miR-26^TKO^*samples at P5, with reasonably low expression in all the W20 samples. The exceptions to this trend were *Wee1,* and *Donson* (DNA replication fork stabilizing factor), which were expressed at higher levels at W20 *miR-26^TKO^*lenses, as well as *Dlgap4,* which was expressed in the P5 *FVB* lenses but peaked in the W20 *miR-26^TKO^* lenses with cataracts.

A previous study explored the function of miR-26 in cultured human lens epithelial cells (SRA01/01) using miR-26 mimics and inhibitory oligonucleotides in an injury-induced anterior subcapsular cataract mouse model^37^. This study suggested that miR-26 inhibits fibrosis by negatively regulating the Jagged-1/Notch signaling pathway. Therefore, differential expression of Jagged-1/Notch signaling pathway genes in the *miR-26^TKO^*datasets was examined (Figure S12). The differentially expressed genes relevant to this pathway generally fell into three patterns of expression (I-III). Genes in group I (including *Numb, Jag2*, and *Hey2*) exhibited very high expression in the W20 *miR-26^TKO^* lenses with cataract, moderate expression in the W20 *miR-26^TKO^* lenses without cataract and low expression in all other conditions. Group II genes (including *Dll1, Tle1,* and *Tle2*) are expressed at low levels in the P5 samples with increased expression in the W20 *FVB* samples and abnormally elevated expression in the W20 *miR-26^TKO^* samples without and with cataract. Group III genes (including *Dll4, Notch 3, Notch4, Tle3, Jag1, Heyl,* and *Hey1*) were expressed at moderate to high levels in the P5 samples with low expression in the W20 *FVB* lenses and abnormally reduced expression in the W20 *miR-26^TKO^*without and with cataracts. These data suggest that expression of genes in the Jagged-Notch signaling pathway are not significantly altered at P5, but by W20, these are either abnormally elevated or reduced in *miR-26^TKO^* lenses. Together, these data suggest that miR-26 is necessary for normal expression of genes in the Jagged-Notch signaling pathway in the lens.

### The role of miR-184 in lens development

To investigate a possible role for *miR-184*, we employed a similar CRISPR/Cas9-based strategy to create a null mutation in this miRNA gene, as described previously (Figure S3). Two guide RNAs (gRNAs) complementary to the template strand of *miR-184* (Figure 8A) were co-injected with Cas9 protein into *FVB/N* zygotes and resultant pups were screened for mutations by PCR and DNA-sequencing. From a total of 70 injected zygotes that were implanted, eight pups were born. Four of these pups contained targeted alleles that were used to generate four independent lines of homozygous *miR-184* KO mice, each of which failed to express mature miR-184-5p transcripts in the lens (Figure 8B). Only one of these lines, *miR-184* KO line 2, was studied in detail. *miR-184* KO line 2 contained a 160 bp deletion in the *miR-184* locus (Figure 8C). In *miR-184* KO line 2, among top 8 potential off-targeted genes (C*blb, Narfl, Ptar1, Slc39a2, Hdac4, Qsox1, Trim3* and *Ipo9*), PCR and sequencing only detected a deletion in *Ptar1* (data not shown) that was eliminated in this line by outcrossing to wild-type *FVB/N* mice.

**Figure 8.**
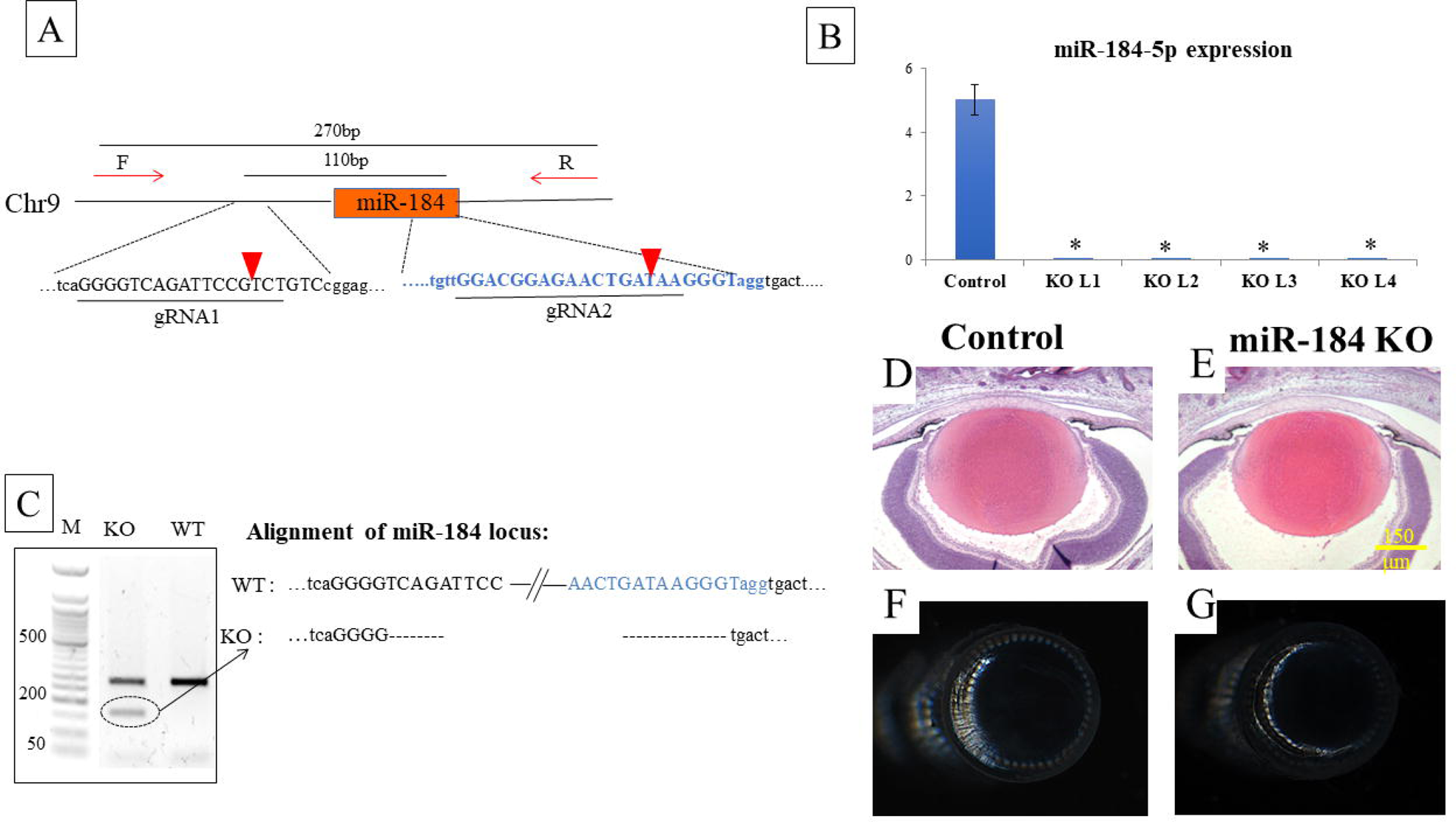
Loss of miR-184 expression did not alter lens morphology. (A) Two gRNAs were targeted 110 bp apart to excise the whole miR-184 sequence. Red arrowheads indicate the cutting sites of the Cas9 enzyme. Indicated primers (red arrows) were used in PCR screening for potential knockout mice. (B) RT-qPCR of 3-week-old lens RNA showed that expression of miR-184-3p was completely abolished in all four *miR-184* KO mouse lines. Error bars on the graph represent SEM. gRNA= guide RNA. (C) PCR screening of DNA from F1 heterozygous mice from the *miR-184* KO line 2 founder (KO) and *FVB/N* (WT) using primers indicated in (A, Table S3), showing the lower band indicating the deleted allele. DNA sequencing of *miR-184* KO line 2 demonstrated a 160 bp deletion. (D-E). Histological analysis of control and miR-184 KO newborn lenses failed to reveal any obvious morphological defect. (F-G) Lenses from 10 month-old *miR-184* KO mice are free of opacity.

Homozygous *miR-184* KO mice all appeared viable and failed to show any obvious phenotype. No histological differences between the control and *miR-184* KO newborn eyes were detected (Figure 8D-E). No opacities or histological abnormalities in eyes from *miR-184 KO* mice were detected when followed up to 10 months of age (Figure 8F-G). To determine if the loss of *miR-184* led to changes in the expression of confirmed and potential miR-184 target genes, we performed RT-qPCR on lenses from 3 week old (P21) control and *miR-184* KO mice (Figure S13). Transcripts from previously identified miR-184 target genes *Ago2*^38^ and *Fzd7*^39^ demonstrated increased expression in *miR-184* KO lenses. In contrast, the level of lens transcripts from the previously identified miR-184 target gene *Numbl* remained unchanged in the *miR-184* KOs, consistent with previous findings demonstrating that miR-184 primarily regulates *Numbl* at the level of translation^40^. The bioinformatics tools TargetScan, and miRWalk were used to identify other potential *miR-184* targets. Five miR-184 targets were predicted by both software tools: Ras-related protein 2A (*Rap2a*), Ras-related protein 2C (*Rap2c*), Lipid phosphate phosphohydrolase 3 (*Ppap2b*), Foxhead box protein O1 (*FoxO1*) and Frizzled 1 (*Fzd1*). Transcripts from all of these predicted targets demonstrated significantly increased expression in *miR-184* KO lenses. Despite these changes in gene expression, we did not detect any overt pathology as examined by microscopy and histology in the *miR-184* knockout mice, suggesting that the loss of miR-184 alone was insufficient to disrupt lens homeostasis in *FVB/N* strain mice at least through the first 10 months of age.

### The role of miR-1 in lens development

The mouse and human genome each contain two copies of the *miR-1* gene (in mice, *miR-1-1* on chromosome 2 and *miR-1-2* on chromosome 18). These genes produce an identical, mature miR-1-3p. The loss of both copies of *miR-1* in mice leads to cardiac failure and perinatal lethality ^41^. We undertook an histological analysis of newborn mouse lenses lacking either *miR-1-1*, *miR-1-2,* or simultaneously both *miR-1-1* and *miR-1-2*. Despite the high expression of *miR-1* in the lens fibers, gross histological analysis of newborn eyes did not reveal any obvious morphological defects in single or double *miR-1* null lenses (Figure S14). While it is possible that *miR-1* deficient mice would exhibit later postnatal lens defects, the lack of a clear newborn phenotype suggests that this miRNA plays no major role in embryonic lens development.

## Discussion

Although we published a comprehensive expression analysis of mRNAs and lncRNAs expressed in the newborn *FVB* mouse lens epithelium and lens fiber cells a decade ago^42^, to our knowledge, no such study has examined the differential expression of miRNAs in these two tissue compartments. As such, this work describing the miRNA expression profile in the lens will serve as a benchmark for evaluating the role of miRNAs in lens development, as well as in pathological conditions. Here, we report the relative abundance and differential expression of miRNAs in the newborn mouse lens epithelium and fiber cells. Of the known miRNAs that we detected in the lens, 184 displayed differential expression between the lens epithelium and lens fiber cells. One of these, miR-1, preferentially expressed in the lens fiber cells, was the fourth most abundantly expressed miR in the newborn mouse lens in our dataset. This result came as a surprise, given a previous report that failed to detect the expression of miR-1 in 4-week old lenses from *C57BL/6* mice by Northern blot^6^. We chose to conduct a functional analysis on three of the four most abundantly expressed miRNAs (miR-184, miR-26, miR-1) in the lens. While they were conducted in whole lenses, other studies also support the high lens expression of these miRNAs^17,22^.

Although embryonic development took place normally in mice lacking all six alleles of *miR-26*, these mice developed bilateral postnatal cataracts between 4 and 22 weeks of age. These cataracts failed to appear in mice lacking only five of the six *miR-26* alleles, attesting to the genetic redundancy of the three *miR-26* genes expressed in the lens. Evaluation of genes differentially expressed between wild-type (*FVB/N*) and *miR-26^TKO^* lenses at 5 days of age pointed to a deregulation of cell proliferation at a stage well before the onset of lens opacities and several key genes linked to cataract (*Aipl1, Ndp, Shh, Otx2, Bfsp2,* and *Myo7a*). Moreover, *miR-26^TKO^*mis-expressed genes also included many candidates linked to cataract as listed in the Cat-Map database, as well as candidates exhibiting lens-enriched expression that are recognized as high-priority in lens biology by the iSyTE database. Further, GSEA analysis of transcripts differentially expressed at 20 weeks of age demonstrated an enrichment for genes associated with complement activation and epithelial to mesenchymal transition. Further, at P5 genes involved in cell proliferation were identified to be mis-expressed, but this effect was not observed at 20 weeks of age. Of the many predicted miR-26 direct target genes, 26 were up-regulated in all *miR-26^TKO^* lens samples as examined analyzed by RNA-seq. These included several immune response genes, including *Lyz2,* encoding a lysozyme; *Lyve1,* encoding a hyaluronan receptor; and *Csf1r,* encoding a receptor that binds both CSF-1 and IL-34. Transcripts for *Slc1a2,* encoding a glutamate transporter, and *Prr5l*, encoding a regulator of mTORC2, were also consistently up-regulated predicted targets in the *miR-26^TKO^*lenses. The up-regulation of these genes suggest that the loss of miR-26 leads to an inflammatory response that is associated with EMT and fibrosis that ultimately leads to cataract.

It is interesting to note that the gene ontology analysis of up-regulated transcripts in the *miR-26^TKO^* samples at 20 weeks identified genes encoding ion channels and other genes important for neuronal development, suggesting a shift in gene expression to that more consistent with neurons. While both lens and nervous system are of ectodermal origin, these two lineages exhibit distinct functional outcomes. Nevertheless, both the lens fiber cells and neurons share several molecular and structural features not commonly found in other tissues, including the expression of nestin, synaptic proteins, glutamate receptors, and GABA receptors^43–47^. Some of these neural characteristics may be driven by molecular regulators of alternative splicing that are shared in both lens and neurons, including several members of the ELAV/Hu proteins^48,49^. Despite this, several recent studies have suggested that the suppression of neural gene expression in lens cells may be an important component of normal lens development and function^50,51^.

A previous study of miR-26 in the human lens epithelial cell line SRA01/04 suggested that miR-26 loss suppressed proliferation and facilitated EMT through the activation of Jagged-1/Notch signaling^37^. This study found that Jag-1 transcripts were directly targeted by miR-26 mimics. However, in our present analysis we could not find any genes associated with Jagged-1/Notch signaling in the common (predicted by both web-based tools) list of the predicted target for miR-26 and DEGs for five day old *miR-26^TKO^* lenses. These findings indicate that the up-regulation of Notch signaling and associated EMT appear as late phenotypes in the *miR-26^TKO^* lenses and could be secondary targets of miR-26.

Our observation not only identifies the possible direct targets of miR-26, but also enhances the use of the lens as a model for EMT as suggested by previous studies^52–56^. Furthermore, the KO of miR-26 shows characteristics such as increased immune response and EMT, resembling PCO and fibrosis. Thus, the relevance of these pathways to known lens pathologies points toward the broad spectrum of possible use of miR-26 as a therapeutic target. Since *miR-26^TKO^*lenses were ruptured at 24-week-old, it is tempting to speculate that this may be due to a combination of factors, including increased lens osmotic pressure, increased EMT, and immune response, which can be examined in the future.

Although we and others have found that miR-204 is expressed abundantly in the lens, we chose not to currently focus on this miRNA given numerous previous investigations of mice in which *miR-204* had been deleted^57–59^. While one of these reports documented adult onset cataracts in mice lacking both *miR-204* and *miR-211*^57^, none of these studies reported congenital cataracts or microphthalmia in mice lacking miR-204 expression, suggesting that embryonic lens development in mice does not require miR-204. Interestingly, a dominant point mutation in *miR-204* is associated with several ocular disorders including early-onset cataracts in humans^60^. In contrast to the apparent normal embryonic lens development in mice lacking *miR-204*, morpholino-induced knockdown of miR-204 in medaka fish disrupted both lens and retina development by interfering with the regulation of Meis2^11^.

Surprisingly, deletion of either of the two abundantly expressed miRNAs (miR-1 and miR-184) had no significant effect on the morphological embryonic development of the mouse lens. miR-1 is most commonly associated with cardiac, skeletal, and smooth muscle development^61–63^, and mice lacking both genomic copies of miR-1 (*miR-1-1* and *miR-1-2*) die from cardiac defects shortly after birth ^41,64^. While the neonatal lethality of *miR-1* KOs prevented analyses of lenses lacking this miRNA beyond birth, we detected no abnormalities in lens size or structure in newborn lenses lacking either or both copies of *miR-1*. Given the lack of obvious developmental abnormalities and the neonatal lethality of the *miR-1* KOs, we did not go beyond histological examination of newborn lenses.

Multiple previous reports have associated a point mutation (+57 C>T) in the seed region of miR-184 associated with human ocular abnormalities, including autosomal dominant severe keratoconus and early onset anterior polar cataract^65–68^, autosomal dominant endothelial dystrophy, iris hypoplasia, congenital cataract, and stromal thinning (EDICT)^65,68^. A recent study found that knocking out miR-184 in zebrafish did not affect embryonic lens development, but these miR-184-deficient zebrafish experienced microphthalmia and cataracts as adults, with no apparent corneal abnormalities^69^. The smaller lens size in these fish was attributed to reduced proliferation and fibrosis that was accompanied by elevated mRNA levels for *cdkn1a* and reduced transcripts for transcription factors *hsf4, ctcf,* and *sox9a*. A previous report of *miR-184* deletion in mice described homozygotes as having elevated levels of TP63 and epidermal hyperplasia^70^. Consistent with our findings, the authors of this study reported that the *miR-184* knockout mice exhibited “no gross phenotype” and were fertile. Given the numerous reports of human ocular abnormalities associated with heterozygous point mutations in *miR-184*^65–68,71^, it was surprising that the homozygous miR-184 knockout mice failed to display any gross developmental or postnatal ocular abnormalities. However, *miR-184* knockout lenses did demonstrate elevated transcript levels for known miR-184 targets: *Ago2* and *Fzd7,* as well as predicted targets *FoxO1, Fzd1, Ppap2b, Rap2a*, and *Rap2c.* The deregulation of these and other genes in the miR-184 knockout mice were not sufficient to disrupt lens morphogenesis or optical clarity, at least on the *FVB/N* genetic background through 10 months of age. It is possible that the dominant ocular phenotypes in human patients with point mutations in miR-184 represent gain of function mutations. Further experiments will be required to clarify the nature of the human ocular abnormalities associated with these miR-184 point mutations.

In summary, the ostensibly normal lens development of *miR-184* KO mice, single/double *miR-26* KO mice, and *miR-1* KO mice in our study are consistent with previous studies showing that deletions of many miRNAs are tolerated due to redundancies between miRNAs and between different pathways. Similarly, less than 10% of miRNA ablations result in developmental defects in *C. elegans*^72^. On the other hand, we provide the first direct evidence that loss of a miRNA family (miR-26) is sufficient to drive cataract formation, and we directly implicate perturbations in inflammation, complement, activation and EMT in driving this phenotype.

## Acknowledgements

We acknowledge and thank the staff (Andor Kiss and Xiaoyun Deng) of the Center for Bioinformatics and Functional Genomics (CBFG) at Miami University for instrumentation and computational support. The authors further thank Peipei Qi, Lin Liu, Aswati Subramanian, and Jacob Weaver for providing their valuable input and/or feedback to the manuscript. We also wish to acknowledge and thank Deepak Srivastava from the Gladstone Institute of Cardiovascular Disease, San Francisco for generously providing tissues from newborn mice lacking either or both *miR-1-1* and *miR 1-2*.

## Data Availability Statement

The sequencing data are available in the Gene Expression Omnibus Database under the following accession number: GSE252611.

## Commercial Relationship Disclosure

No commercial relationships to disclose

---

**Figure S1. miRNA-seq on newborn lens epithelium and fibers.** Distance matrix indicates concurrence in clustering between sample types.

**Figure S2. RT-qPCR analysis of miR-1, miR-184 and miR-26 expression in lens epithelium and lens fibers.** The expression of miR-1-3p in lens epithelium and lens fibers normalized SnoRNA202 (A) or GAPDH (B). The expression of miR-184-3p (C), miR-26a-5p (D), and miR-26-5p (E) was normalized to SnoRNA202 by RT-qPCR. Only miR-1-3p exhibited significantly enriched expression in lens fibers. N.S. = not significant.

**Figure S3. Generalized strategy for targeting miR-loci for CRISPR/Cas9 mediated deletion.** Two specific guide RNAs (gRNAs) flanking the targeted miRNA sequence were preincubated with Cas9 protein prior to microinjection into *FVB/N* zygotes. Injected zygotes were transferred to pseudopregnant mice and resultant pups were screened for the targeted deletion by PCR.

**Figure S4. Deletion of miR-26 family members did not affect expression of their host genes.** Deletion of *miR-26a1*, *miR-26a2,* and *miR-26b* genomic sequences did not significantly affect the relative expression of their host genes. Error bars on the graph represent SEM. N.S = no significant difference.

**Figure S5. Cataract-associated genes that are up-regulated in the *miR-26^TKO^* lenses.** (A) Venn diagram displays the intersections of genes that are up-regulated in *miR-26^TKO^* lenses at P5 and W20 with and without cataract with genes in the Cat-Map list. (B) Heatmap using z-score adjusted expression values all genes in the Cat-Map list that are up-regulated in any of the *miR-26^TKO^* lens samples.

**Figure S6. Cataract-associated genes that are down-regulated in the *miR-26^TKO^* lenses.** (A) Venn diagram displays the intersections of genes that are down-regulated in *miR-26^TKO^* lenses at P5 and W20 with and without cataract with genes in the Cat-Map list. (B) Heatmap using z-score adjusted expression values all genes in the Cat-Map list that are down-regulated in any of the *miR-26^TKO^* lens samples.

**Figure S7. iSyTE genes that are up-regulated in the *miR-26^TKO^*lenses.** (A) Venn diagram displays the intersections of genes that are up-regulated in *miR-26^TKO^* lenses at P5 and W20 with and without cataract with genes listed as lens-enriched in iSyTE. (B) Heatmap using z-score adjusted expression values all genes in the lens-enriched iSyTE list that are up-regulated in any of the *miR-26^TKO^*lens samples.

**Figure S8. iSyTE genes that are down-regulated in the *miR-26^TKO^*lenses.** (A) Venn diagram displays the intersections of genes that are down-regulated in *miR-26^TKO^* lenses at P5 and W20 with and without cataract with genes listed as lens-enriched in iSyTE. (B) Heatmap using z-score adjusted expression values all genes in the lens-enriched iSyTE list that are down-regulated in any of the *miR-26^TKO^*lens samples.

**Figure S9, Venn diagram shows the predicted gene targets by miRwalk (blue) and Targetscan (Yellow).** Common targets predicted by both web-based software tools were used for further analysis.

AFigure S10. Venn diagram represents the genes down-regulated in *miR-26^TKO^*lenses compared to *FVB* control lenses. The down-regulated genes were intersected with predicted targets for miR-26.

**Figure S11. Gene set enrichment analysis identifies proliferation enrichment in *miR-26^TKO^* mice at P5 stage**. (A-B) GSEA enrichment plot represents E2F targets and G2M checkpoint in *miR-26* TKO_P5 stage (red) while compared with P5 *FVB/N* mice (blue). Genes are ordered along the x-axis based on expression rank between the two conditions. Black bars indicated genes associated with the given term. Green line indicates the enrichment score determined by GSEA. (C) Heatmap displays expression values scaled by z-score revealing highest expression of E2F target genes in the TKO_P5 samples relative to other conditions.

**Figure S12. Expression analysis of genes related to Notch-signaling in the *miR-26^TKO^* lenses.** Heatmap displays expression values scaled by z-score revealing three categories of expression patterns of Jagged-1/Notch signaling genes.

**Figure S13. Loss of miR-1 expression did not alter newborn lens morphology.** Histological staining of newborn mouse eye sections from control (A), single deletion (B and C) and double deletions (D) of *miR-1* genes. Gross histological analysis did not reveal any obvious defects.

**Figure S14. Gene expression changes in miR-184 KO lenses.** Relative expression of putative (black letters) and known miR-184-targeted genes (red letters) in 3-week-old lenses from control and *miR-184* KO lenses was quantified by RT-qPCR. Error bars on the graph represent SEM and the asterisk represents a significant difference from the control value. N.S, no significance.

## References

1. Cvekl A, Zhang X. Signaling and Gene Regulatory Networks in Mammalian Lens Development. Trends Genet. 2017;33(10):677–702.

2. Lachke SA. RNA-binding proteins and post-transcriptional regulation in lens biology and cataract: Mediating spatiotemporal expression of key factors that control the cell cycle, transcription, cytoskeleton and transparency. Exp Eye Res. 2022;214:108889.

3. Li Y, Piatigorsky J. Targeted deletion of Dicer disrupts lens morphogenesis, corneal epithelium stratification, and whole eye development. Dev Dyn. 2009;238(9):2388–2400.

4. Shaham O, Gueta K, Mor E, et al. Pax6 regulates gene expression in the vertebrate lens through miR-204. PLoS Genet. 2013;9(3):e1003357.

5. Chen S, Zhang C, Shen L, Hu J, Chen X, Yu Y. Noncoding RNAs in cataract formation: Star molecules emerge in an endless stream. Pharmacol Res. 2022;184:106417.

6. Frederikse PH, Donnelly R, Partyka LM. miRNA and Dicer in the mammalian lens: expression of brain-specific miRNAs in the lens. Histochem Cell Biol. 2006;126(1):1–8.

7. Karali M, Peluso I, Gennarino VA, et al. miRNeye: a microRNA expression atlas of the mouse eye. BMC Genomics. 2010;11(1):1–14.

8. Karali M, Peluso I, Marigo V, Banfi S. Identification and Characterization of MicroRNAs Expressed in the Mouse Eye. Invest Ophthalmol Vis Sci. 2007;48(2):509–515.

9. Ryan DG, Oliveira-Fernandes M, Lavker RM. MicroRNAs of the mammalian eye display distinct and overlapping tissue specificity. Mol Vis. 2006;12:1175–1184.

10. Wolf L, Gao CS, Gueta K, et al. Identification and characterization of FGF2-dependent mRNA: microRNA networks during lens fiber cell differentiation. G3. 2013;3(12):2239–2255.

11. Conte I, Carrella S, Avellino R, et al. miR-204 is required for lens and retinal development via Meis2 targeting. Proc Natl Acad Sci U S A. 2010;107(35):15491–15496.

12. Wurm A, Sock E, Fuchshofer R, Wegner M, Tamm ER. Anterior segment dysgenesis in the eyes of mice deficient for the high-mobility-group transcription factor Sox11. Exp Eye Res. 2008;86(6):895–907.

13. Pillai-Kastoori L, Wen W, Wilson SG, et al. Sox11 is required to maintain proper levels of Hedgehog signaling during vertebrate ocular morphogenesis. PLoS Genet. 2014;10(7):e1004491.

14. Shi Y, Tu Y, Mecham RP, Bassnett S. Ocular phenotype of Fbn2-null mice. Invest Ophthalmol Vis Sci. 2013;54(12):7163–7173.

15. Wang Y, Li W, Zang X, et al. MicroRNA-204-5p regulates epithelial-to-mesenchymal transition during human posterior capsule opacification by targeting SMAD4. Invest Ophthalmol Vis Sci. 2013;54(1):323–332.

16. Hoffmann A, Huang Y, Suetsugu-Maki R, et al. Implication of the miR-184 and miR-204 competitive RNA network in control of mouse secondary cataract. Mol Med. 2012;18(3):528–538.

17. Anand D, Al Saai S, Shrestha SK, Barnum CE, Chuma S, Lachke SA. Genome-Wide Analysis of Differentially Expressed miRNAs and Their Associated Regulatory Networks in Lenses Deficient for the Congenital Cataract-Linked Tudor Domain Containing Protein TDRD7. Front Cell Dev Biol. 2021;9:615761.

18. Peng CH, Liu JH, Woung LC, et al. MicroRNAs and cataracts: correlation among let-7 expression, age and the severity of lens opacity. Br J Ophthalmol. 2012;96(5):747–751.

19. Chien KH, Chen SJ, Liu JH, et al. Correlation between microRNA-34a levels and lens opacity severity in age-related cataracts. EYE. 2013;27(7):883–888.

20. Wu C, Lin H, Wang Q, et al. Discrepant expression of microRNAs in transparent and cataractous human lenses. Invest Ophthalmol Vis Sci. 2012;53(7):3906–3912.

21. Qin Y, Zhao J, Min X, et al. MicroRNA-125b inhibits lens epithelial cell apoptosis by targeting p53 in age-related cataract. Biochim Biophys Acta. 2014;1842(12 Pt A):2439–2447.

22. Khan SY, Hackett SF, Lee MCW, Pourmand N, Conover Talbot C, Amer Riazuddin S. Transcriptome Profiling of Developing Murine Lens Through RNA Sequencing. Invest Ophthalmol Vis Sci. 2015;56(8):4919–4926.

23. Tian L, Huang K, DuHadaway JB, Prendergast GC, Stambolian D. Genomic profiling of miRNAs in two human lens cell lines. Curr Eye Res. 2010;35(9):812–818.

24. Kim D, Paggi JM, Park C, Bennett C, Salzberg SL. Graph-based genome alignment and genotyping with HISAT2 and HISAT-genotype. Nat Biotechnol. 2019;37(8):907–915.

25. Pertea M, Pertea GM, Antonescu CM, Chang TC, Mendell JT, Salzberg SL. StringTie enables improved reconstruction of a transcriptome from RNA-seq reads. Nat Biotechnol. 2015;33(3):290–295.

26. Love MI, Huber W, Anders S. Moderated estimation of fold change and dispersion for RNA-seq data with DESeq2. Genome Biol. 2014;15(12):550.

27. Friedländer MR, Mackowiak SD, Li N, Chen W, Rajewsky N. miRDeep2 accurately identifies known and hundreds of novel microRNA genes in seven animal clades. Nucleic Acids Res. 2012;40(1):37–52.

28. Kolberg L, Raudvere U, Kuzmin I, Adler P, Vilo J, Peterson H. g:Profiler-interoperable web service for functional enrichment analysis and gene identifier mapping (2023 update). Nucleic Acids Res. 2023;51(W1):W207–W212.

29. Sherman BT, Hao M, Qiu J, et al. DAVID: a web server for functional enrichment analysis and functional annotation of gene lists (2021 update). Nucleic Acids Res. 2022;50(W1):W216–W221.

30. Huang DW, Sherman BT, Lempicki RA. Systematic and integrative analysis of large gene lists using DAVID bioinformatics resources. Nat Protoc. 2009;4(1):44–57.

31. Oliveros JC. Venny. An interactive tool for comparing lists with Venn’s diagrams. Venny 2.1. Published 2007-2015. Accessed November 10, 2022. https://bioinfogp.cnb.csic.es/tools/venny/index.html

32. Sticht C, De La Torre C, Parveen A, Gretz N. miRWalk: An online resource for prediction of microRNA binding sites. PLoS One. 2018;13(10):e0206239.

33. McGeary SE, Lin KS, Shi CY, et al. The biochemical basis of microRNA targeting efficacy. Science. 2019;366(6472). doi:10.1126/science.aav1741

34. Subramanian A, Tamayo P, Mootha VK, et al. Gene set enrichment analysis: a knowledge-based approach for interpreting genome-wide expression profiles. Proc Natl Acad Sci U S A. 2005;102(43):15545–15550.

35. Shiels A, Bennett TM, Hejtmancik JF. Cat-Map: putting cataract on the map. Mol Vis. 2010;16:2007–2015.

36. Kakrana A, Yang A, Anand D, et al. iSyTE 2.0: a database for expression-based gene discovery in the eye. Nucleic Acids Res. 2018;46(D1):D875–D885.

37. Chen X, Xiao W, Chen W, et al. MicroRNA-26a and -26b inhibit lens fibrosis and cataract by negatively regulating Jagged-1/Notch signaling pathway. Cell Death Differ. 2017;24(8):1431–1442.

38. Tattikota SG, Rathjen T, McAnulty SJ, et al. Argonaute2 mediates compensatory expansion of the pancreatic β cell. Cell Metab. 2014;19(1):122–134.

39. Takahashi Y, Chen Q, Rajala RVS, Ma JX. MicroRNA-184 modulates canonical Wnt signaling through the regulation of frizzled-7 expression in the retina with ischemia-induced neovascularization. FEBS Lett. 2015;589(10):1143–1149.

40. Liu C, Teng ZQ, Santistevan NJ, et al. Epigenetic regulation of miR-184 by MBD1 governs neural stem cell proliferation and differentiation. Cell Stem Cell. 2010;6(5):433–444.

41. Heidersbach A, Saxby C, Carver-Moore K, et al. microRNA-1 regulates sarcomere formation and suppresses smooth muscle gene expression in the mammalian heart. Elife. 2013;2:e01323.

42. Hoang TV, Kumar PKR, Sutharzan S, Tsonis PA, Liang C, Robinson ML. Comparative transcriptome analysis of epithelial and fiber cells in newborn mouse lenses with RNA sequencing. Mol Vis. 2014;20:1491–1517.

43. Frederikse PH, Kasinathan C, Kleiman NJ. Parallels between neuron and lens fiber cell structure and molecular regulatory networks. Dev Biol. 2012;368(2):255–260.

44. Frederikse PH, Nandanoor A, Kasinathan C. “Moonlighting” GAPDH Protein Localizes with AMPA Receptor GluA2 and L1 Axonal Cell Adhesion Molecule at Fiber Cell Borders in the Lens. Curr Eye Res. 2016;41(1):41–49.

45. Frederikse PH, Nandanoor A, Kasinathan C. Fragile X Syndrome FMRP Co-localizes with Regulatory Targets PSD-95, GABA Receptors, CaMKIIα, and mGluR5 at Fiber Cell Membranes in the Eye Lens. Neurochem Res. 2015;40(11):2167–2176.

46. Frederikse PH, Kasinathan C. Lens GABA receptors are a target of GABA-related agonists that mitigate experimental myopia. Med Hypotheses. 2015;84(6):589–592.

47. Yang J, Bian W, Gao X, Chen L, Jing N. Nestin expression during mouse eye and lens development. Mech Dev. 2000;94(1-2):287–291.

48. Bitel CL, Perrone-Bizzozero NI, Frederikse PH. HuB/C/D, nPTB, REST4, and miR-124 regulators of neuronal cell identity are also utilized in the lens. Mol Vis. 2010;16:2301–2316.

49. Hilgers V. Regulation of neuronal RNA signatures by ELAV/Hu proteins. Wiley Interdiscip Rev RNA. 2023;14(2):e1733.

50. Maddala R, Gao J, Mathias RT, et al. Absence of S100A4 in the mouse lens induces an aberrant retina-specific differentiation program and cataract. Sci Rep. 2021;11(1):2203.

51. Tangeman JA, Rebull SM, Grajales-Esquivel E, et al. Integrated single-cell multiomics uncovers foundational regulatory mechanisms of lens development and pathology. Development. 2024;151(1). doi:10.1242/dev.202249

52. Eldred JA, Dawes LJ, Wormstone IM. The lens as a model for fibrotic disease. Philos Trans R Soc Lond B Biol Sci. 2011;366(1568):1301–1319.

53. Taiyab A, West-Mays J. Lens Fibrosis: Understanding the Dynamics of Cell Adhesion Signaling in Lens Epithelial-Mesenchymal Transition. Front Cell Dev Biol. 2022;10:886053.

54. Upreti A, Padula SL, Tangeman JA, et al. Lens Epithelial Explants Treated with Vitreous Humor Undergo Alterations in Chromatin Landscape with Concurrent Activation of Genes Associated with Fiber Cell Differentiation and Innate Immune Response. Cells. 2023;12(3). doi:10.3390/cells12030501

55. Lovicu FJ, Shin EH, McAvoy JW. Fibrosis in the lens. Sprouty regulation of TGFβ-signaling prevents lens EMT leading to cataract. Exp Eye Res. 2016;142:92–101.

56. Shu DY, Lovicu FJ. Enhanced EGF receptor-signaling potentiates TGFβ-induced lens epithelial-mesenchymal transition. Exp Eye Res. 2019;185:107693.

57. Huang J, Zhao L, Fan Y, et al. The microRNAs miR-204 and miR-211 maintain joint homeostasis and protect against osteoarthritis progression. Nat Commun. 2019;10(1):2876.

58. Jo S, Chen J, Xu G, Grayson TB, Thielen LA, Shalev A. miR-204 Controls Glucagon-Like Peptide 1 Receptor Expression and Agonist Function. Diabetes. 2018;67(2):256–264.

59. Cheng Y, Wang D, Wang F, et al. Endogenous miR-204 Protects the Kidney against Chronic Injury in Hypertension and Diabetes. J Am Soc Nephrol. 2020;31(7):1539–1554.

60. Jedlickova J, Vajter M, Barta T, et al. MIR204 n.37C>T variant as a cause of chorioretinal dystrophy variably associated with iris coloboma, early-onset cataracts and congenital glaucoma. Clin Genet. 2023;104(4):418–426.

61. Zhao Y, Samal E, Srivastava D. Serum response factor regulates a muscle-specific microRNA that targets Hand2 during cardiogenesis. Nature. 2005;436(7048):214–220.

62. Chen JF, Mandel EM, Thomson JM, et al. The role of microRNA-1 and microRNA-133 in skeletal muscle proliferation and differentiation. Nat Genet. 2006;38(2):228–233.

63. Xie C, Huang H, Sun X, et al. MicroRNA-1 regulates smooth muscle cell differentiation by repressing Kruppel-like factor 4. Stem Cells Dev. 2011;20(2):205–210.

64. Zhao Y, Ransom JF, Li A, et al. Dysregulation of cardiogenesis, cardiac conduction, and cell cycle in mice lacking miRNA-1-2. Cell. 2007;129(2):303–317.

65. Hughes AE, Bradley DT, Campbell M, et al. Mutation altering the miR-184 seed region causes familial keratoconus with cataract. Am J Hum Genet. 2011;89(5):628–633.

66. Bykhovskaya Y, Caiado Canedo AL, Wright KW, Rabinowitz YS. C.57 C > T Mutation in MIR 184 is Responsible for Congenital Cataracts and Corneal Abnormalities in a Five-generation Family from Galicia, Spain. Ophthalmic Genet. 2015;36(3):244–247.

67. Farzadfard A, Nassiri N, Moghadam TN, Paylakhi SH, Elahi E. Screening for MIR184 Mutations in Iranian Patients with Keratoconus. J Ophthalmic Vis Res. 2016;11(1):3–7.

68. Iliff BW, Riazuddin SA, Gottsch JD. A single-base substitution in the seed region of miR-184 causes EDICT syndrome. Invest Ophthalmol Vis Sci. 2012;53(1):348–353.

69. Zhang J, Li P, Sun L, et al. Knockout of miR-184 in zebrafish leads to ocular abnormalities by elevating p21 levels. FASEB J. 2023;37(5):e22927.

70. Nagosa S, Leesch F, Putin D, et al. microRNA-184 Induces a Commitment Switch to Epidermal Differentiation. Stem Cell Reports. 2017;9(6):1991–2004.

71. Lechner J, Bae HA, Guduric-Fuchs J, et al. Mutational analysis of MIR184 in sporadic keratoconus and myopia. Invest Ophthalmol Vis Sci. 2013;54(8):5266–5272.

72. Alvarez-Saavedra E, Horvitz HR. Many families of C. elegans microRNAs are not essential for development or viability. Curr Biol. 2010;20(4):367–373.

