## Supplementary figures and images for "miR-26 deficiency causes alterations in lens transcriptome and results in adult-onset cataract"

### Figure S1.tif

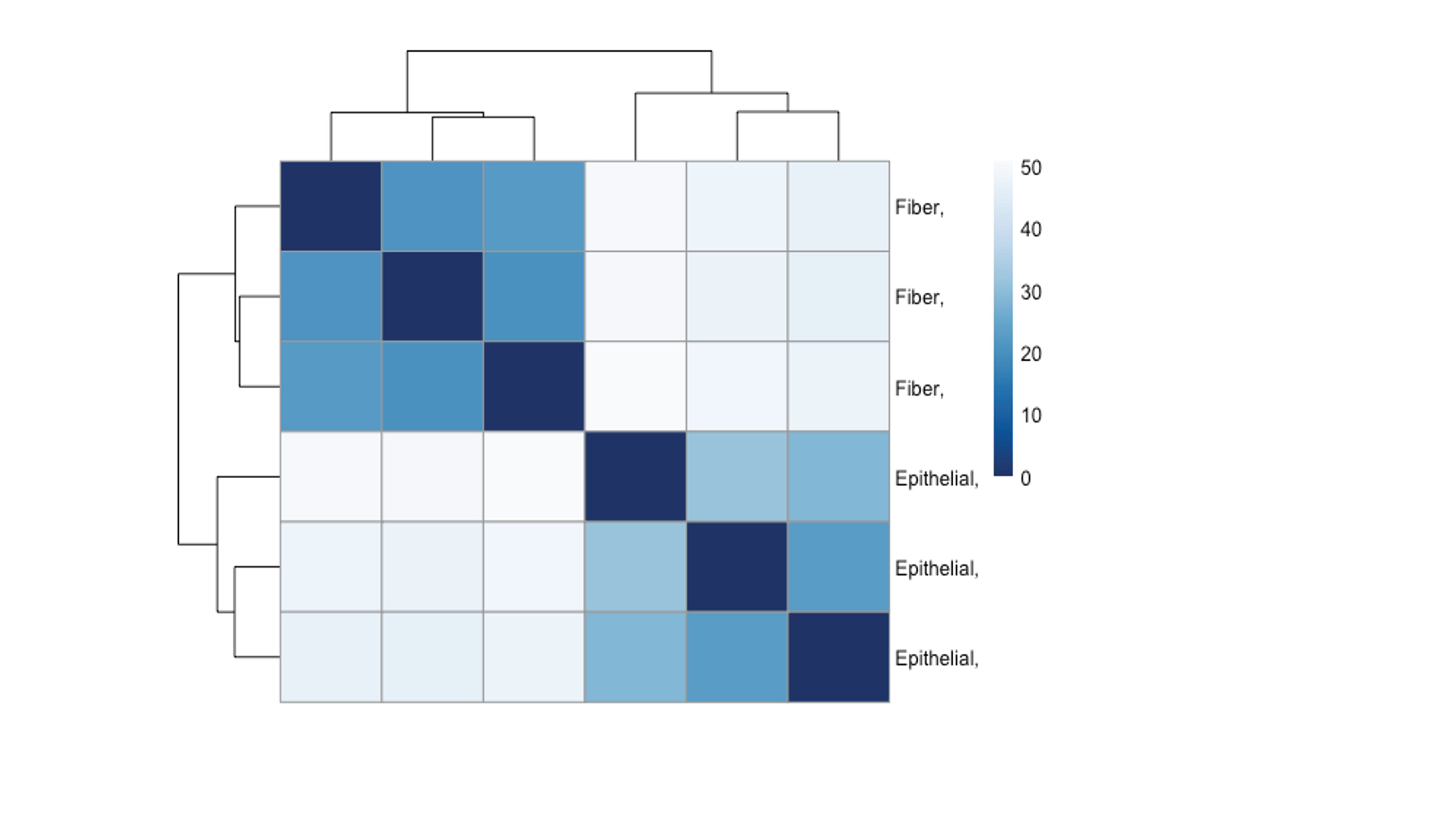

### Figure S2.tif

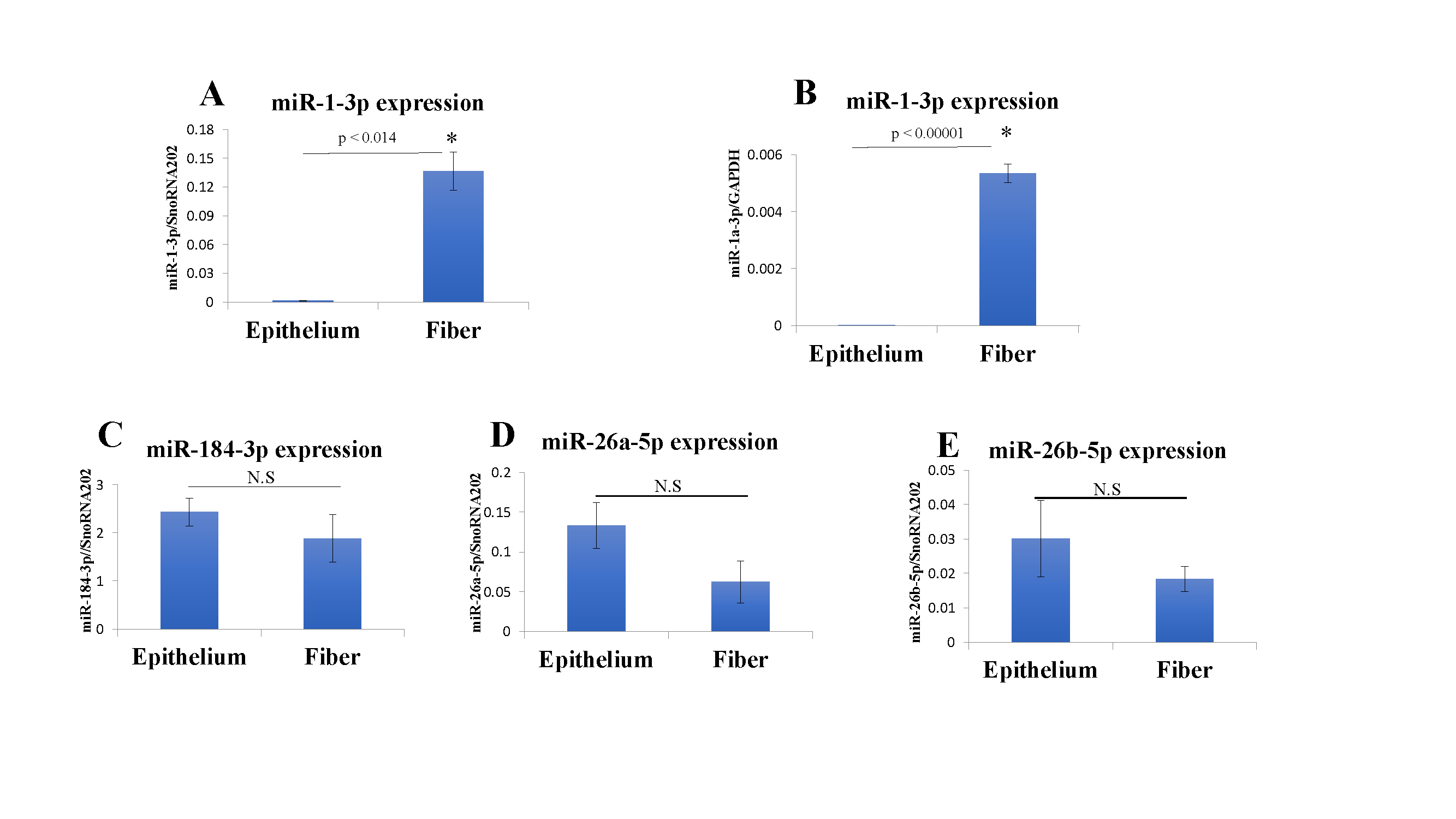

### Figure S3.tif

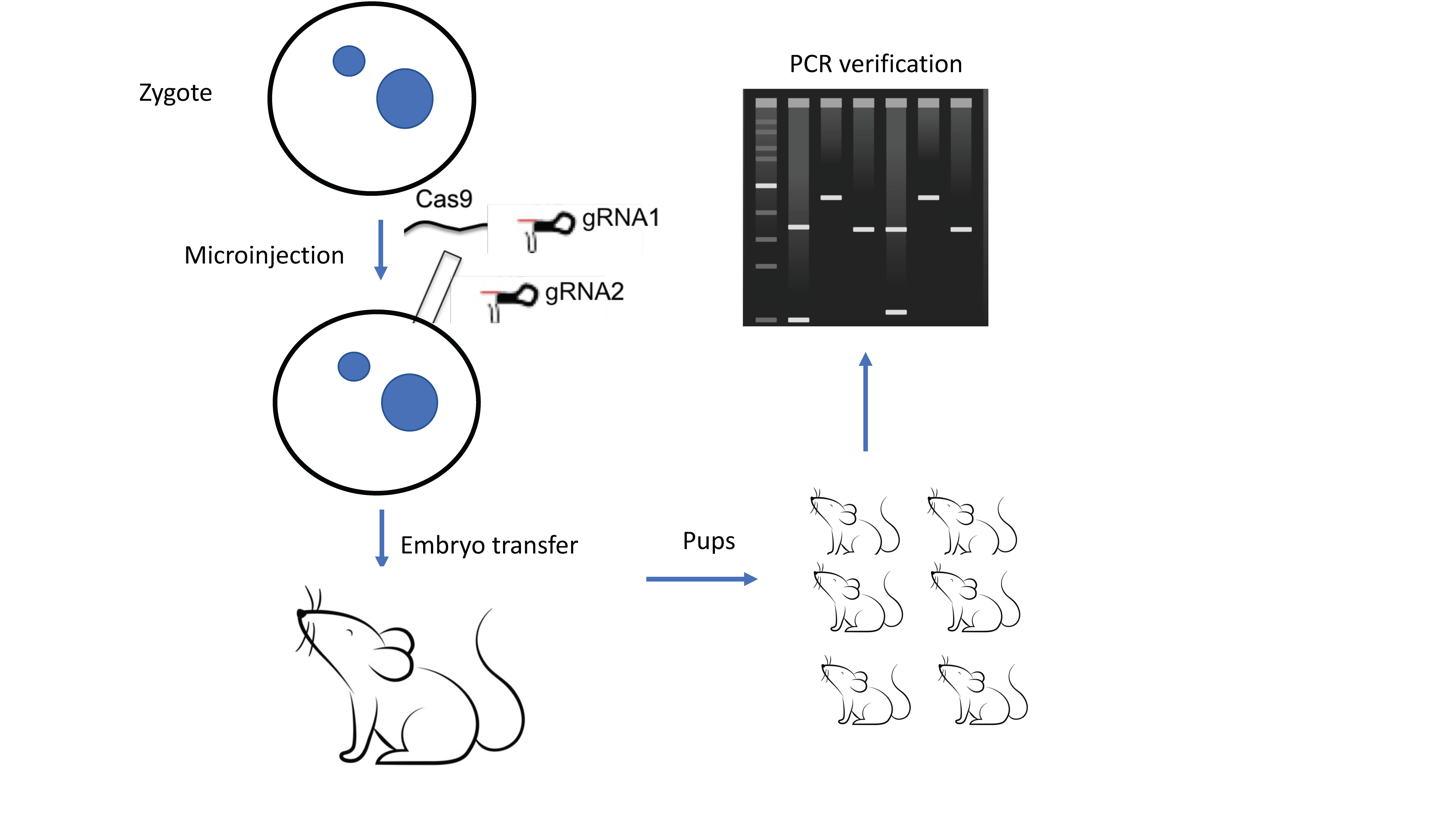

### Figure S4.tif

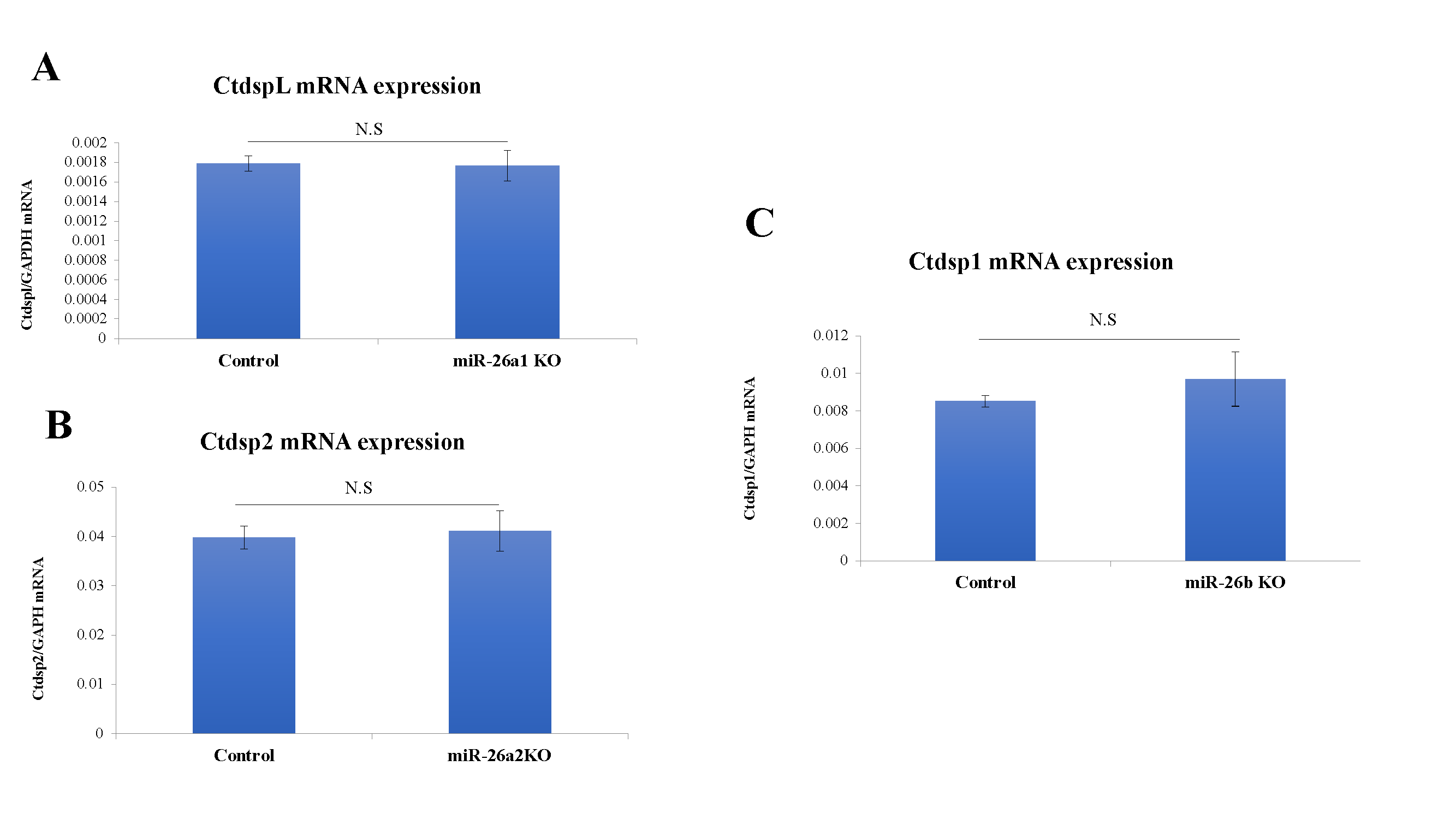

### Figure S5.tif

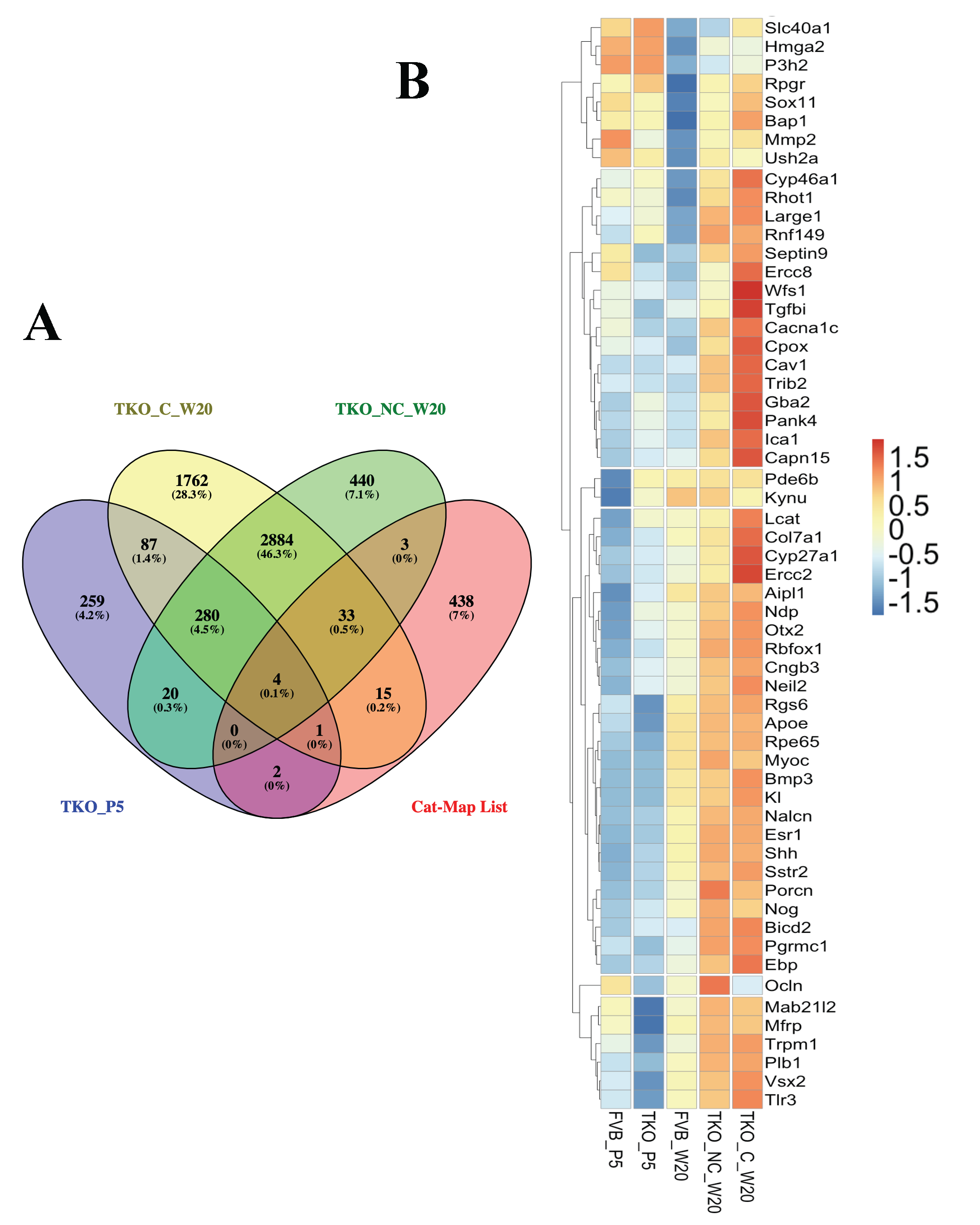

### Figure S6.tif

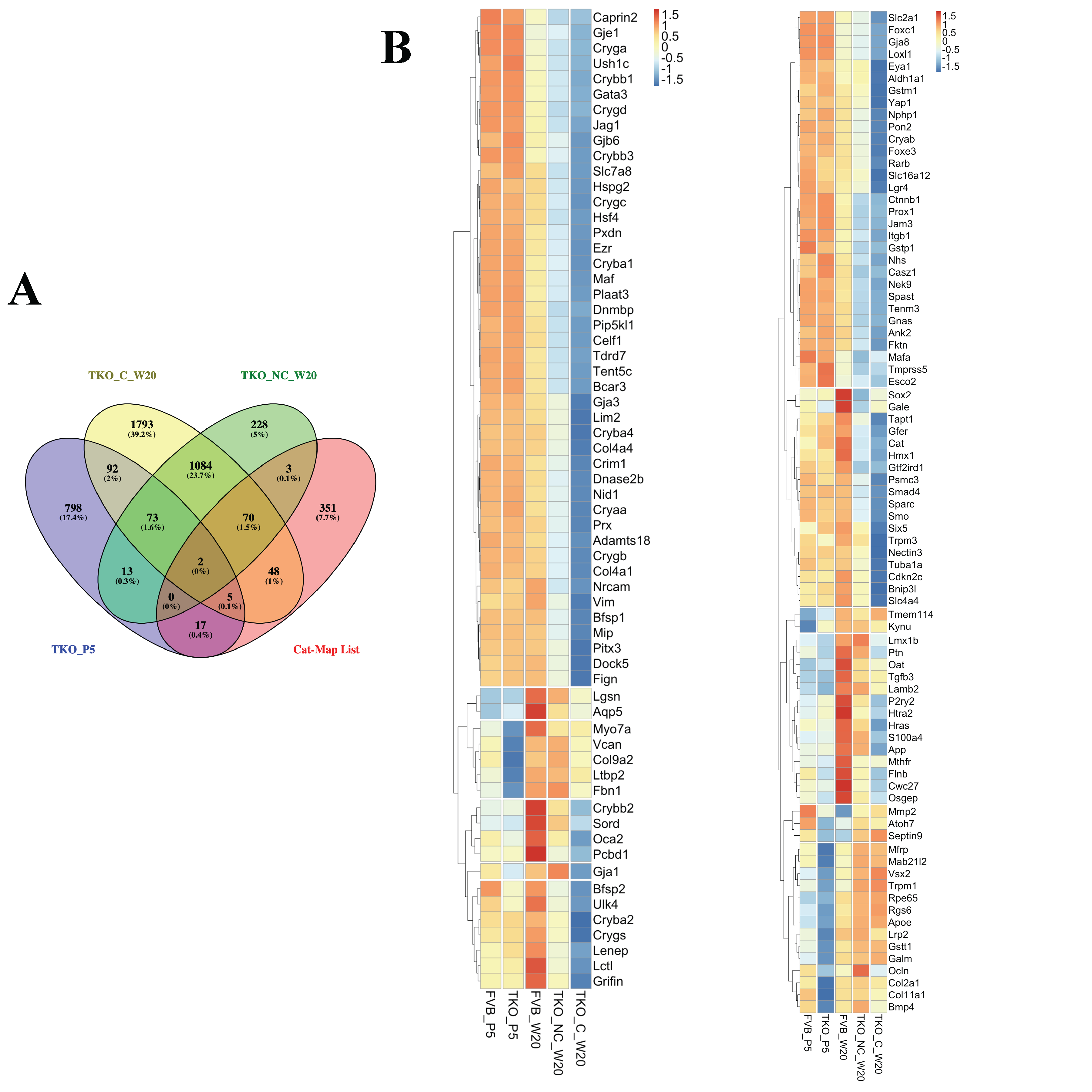

### Figure S7.tif

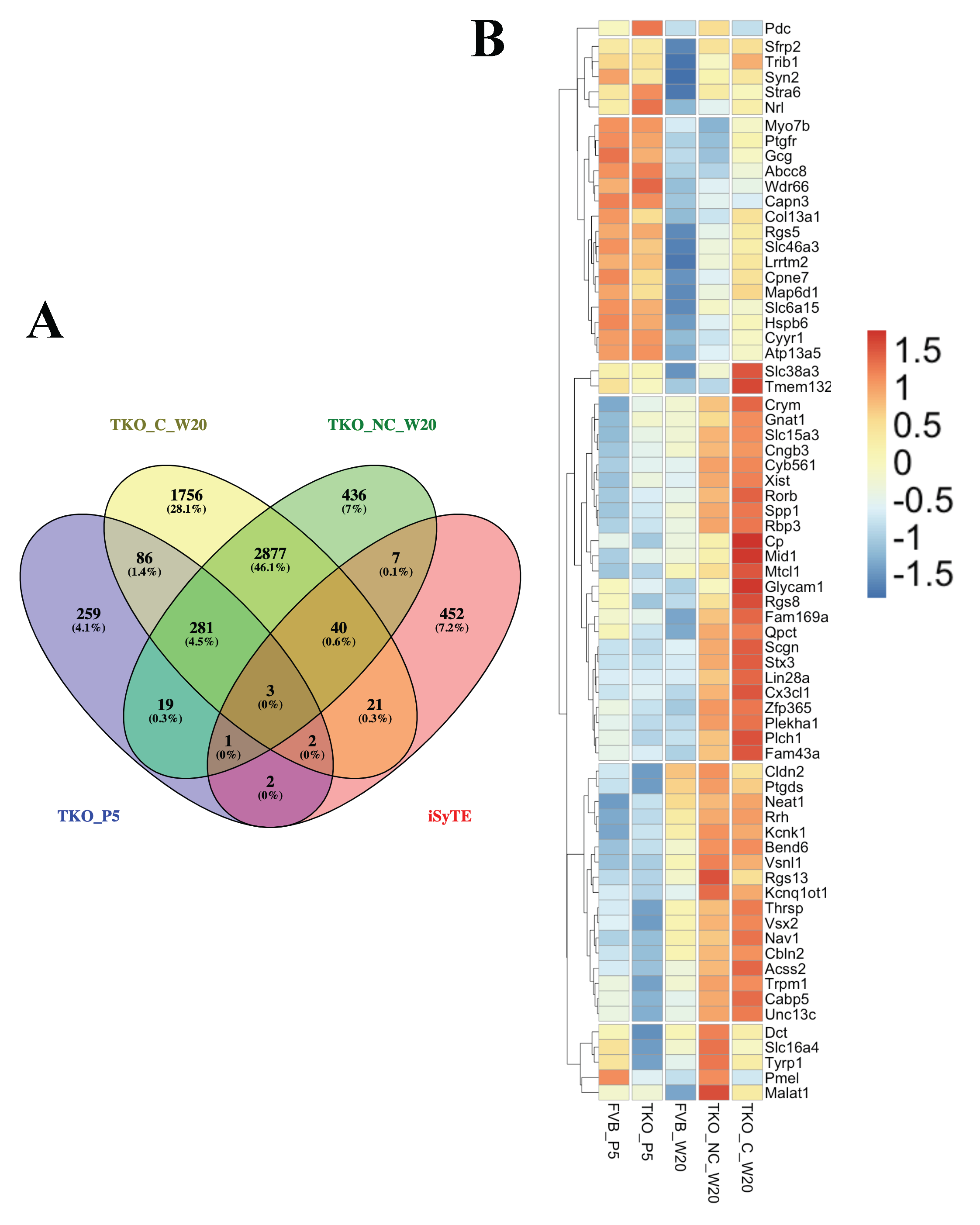

### Figure S8.tif

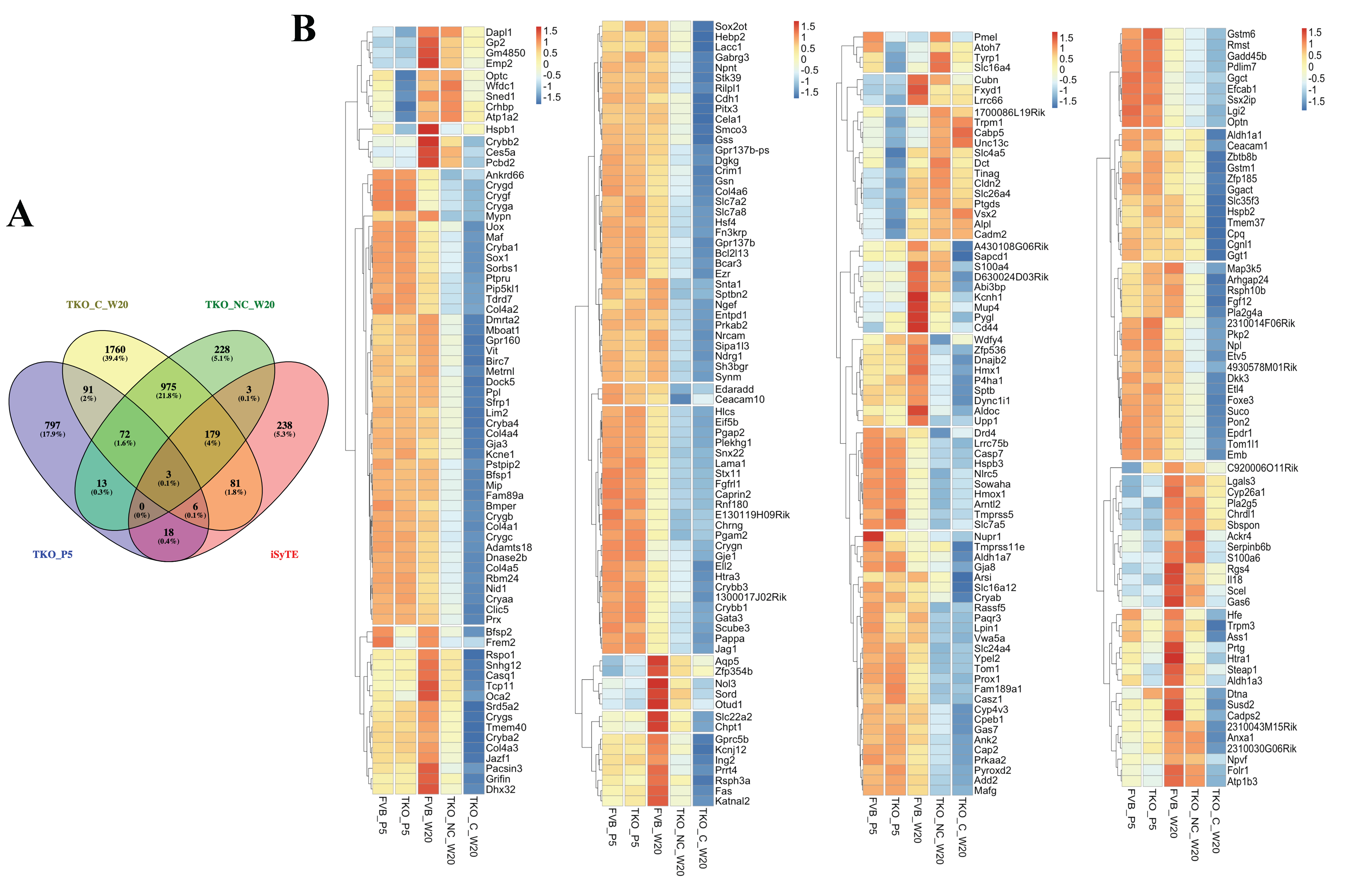

### Figure S9.tif

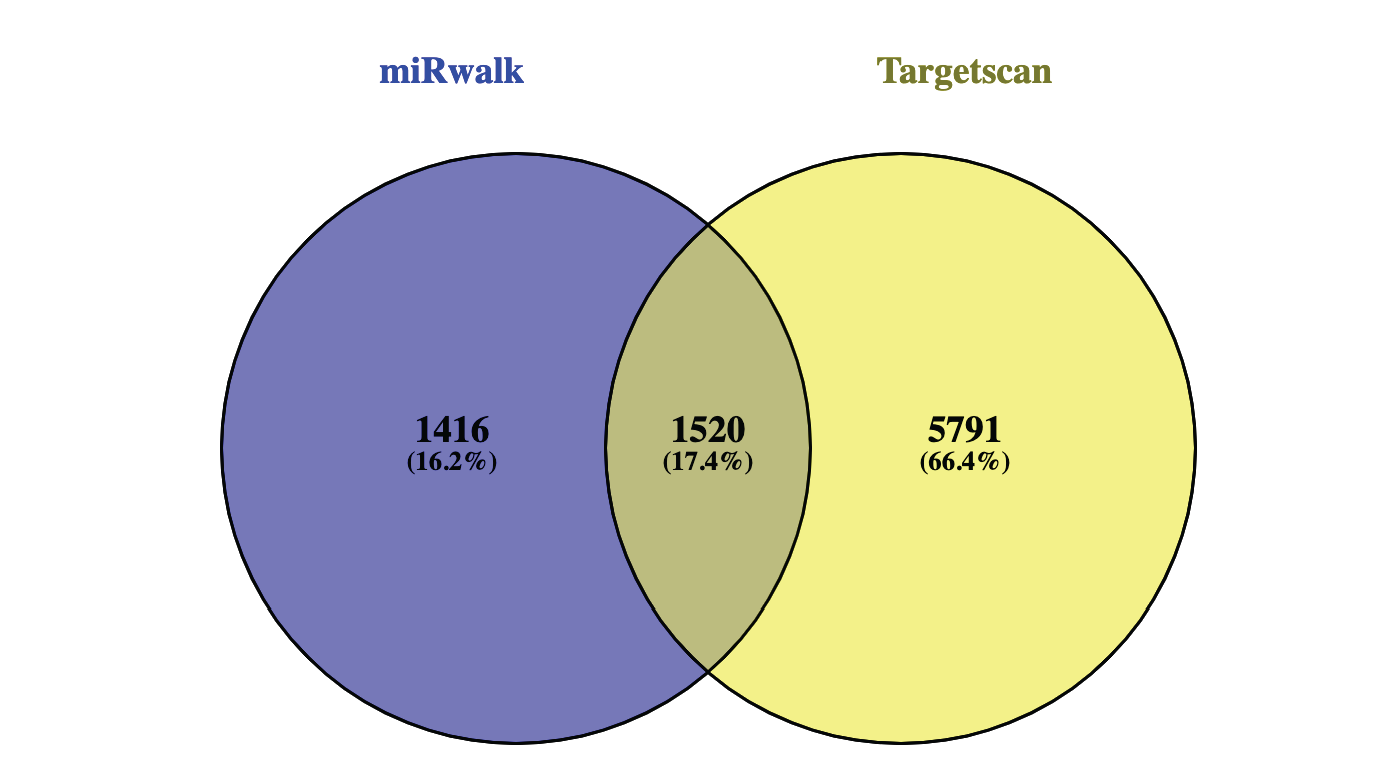

### Figure S10.tif

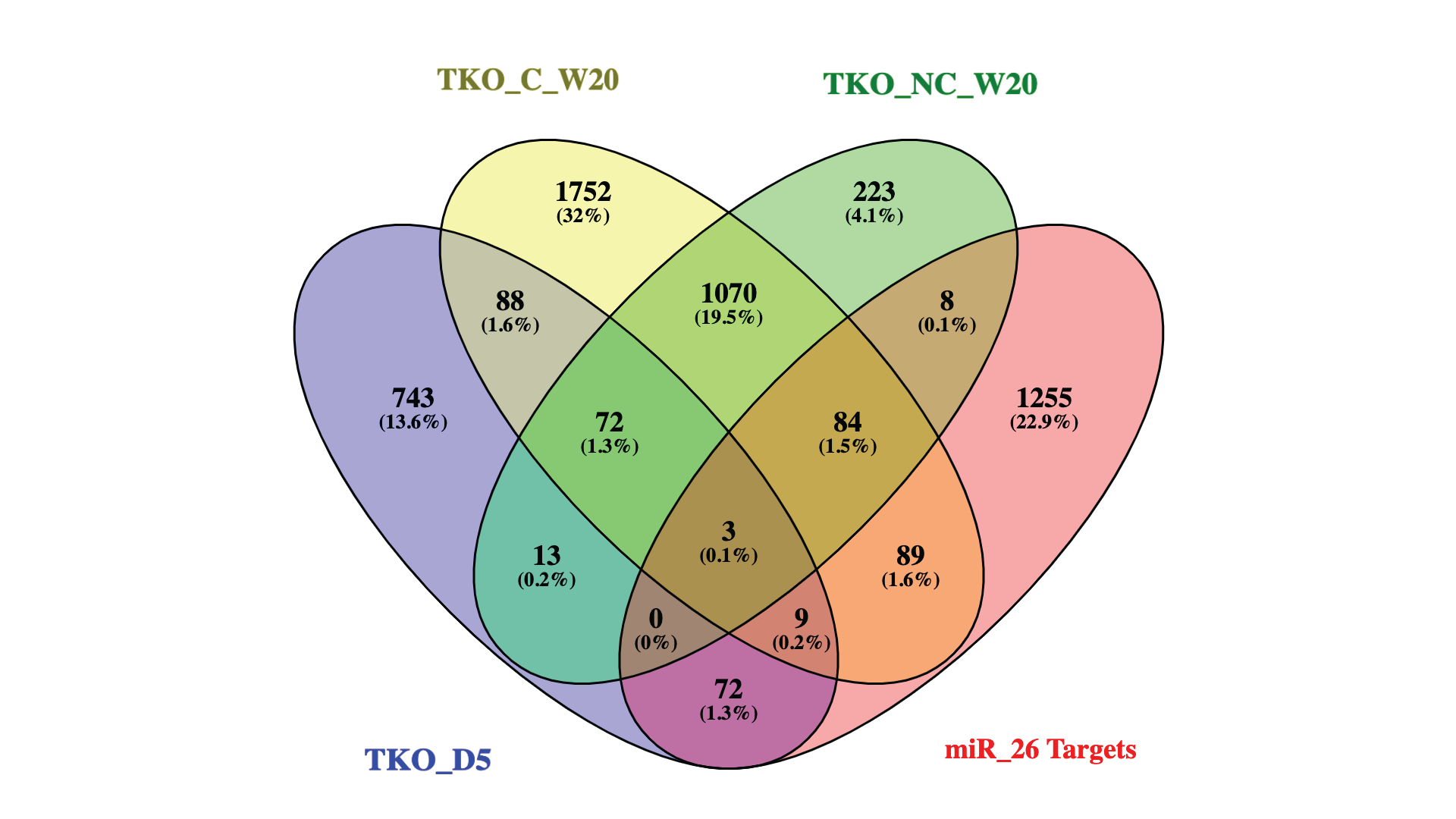

### Figure S11.tif

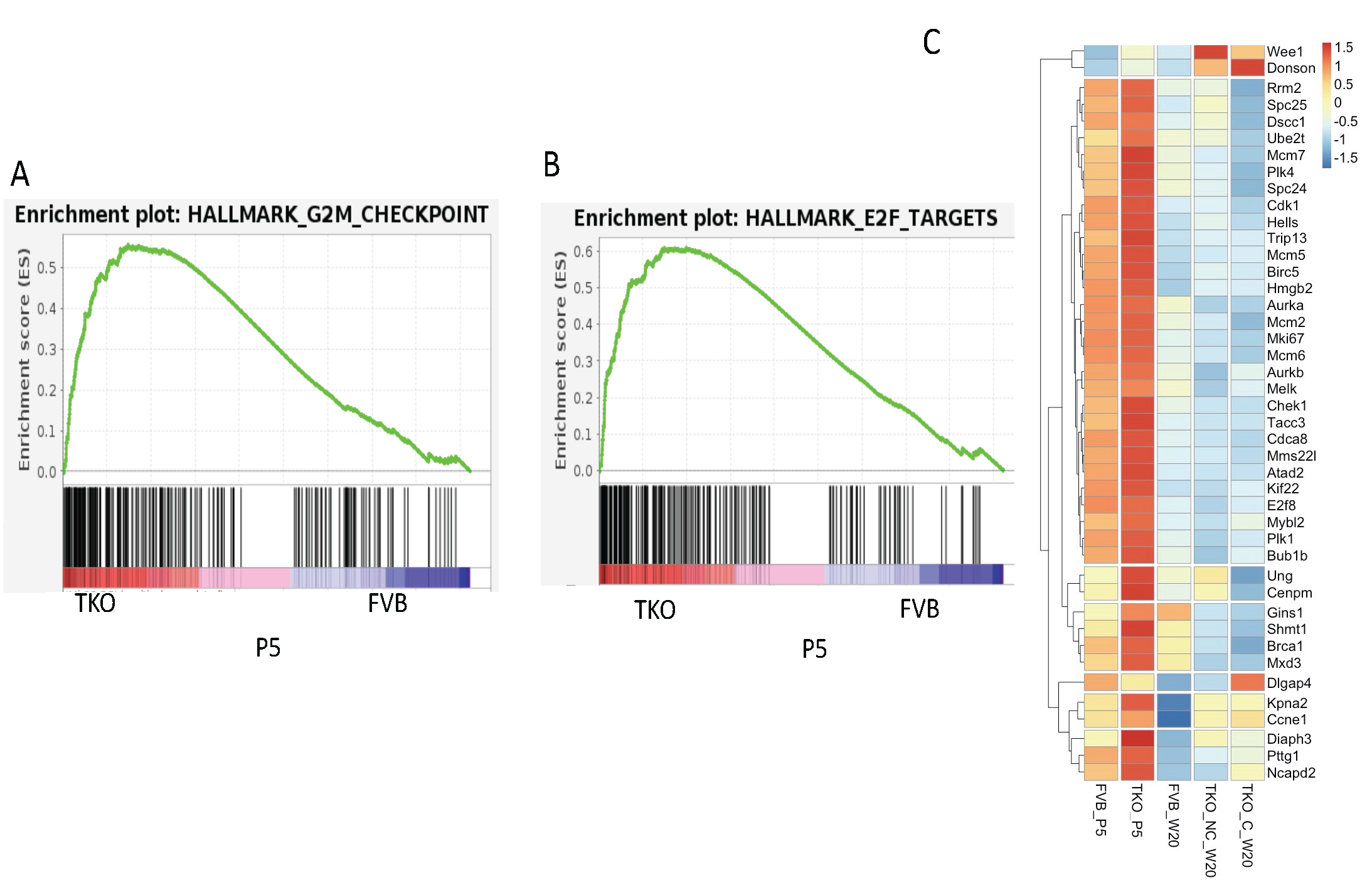

### Figure S12.tif

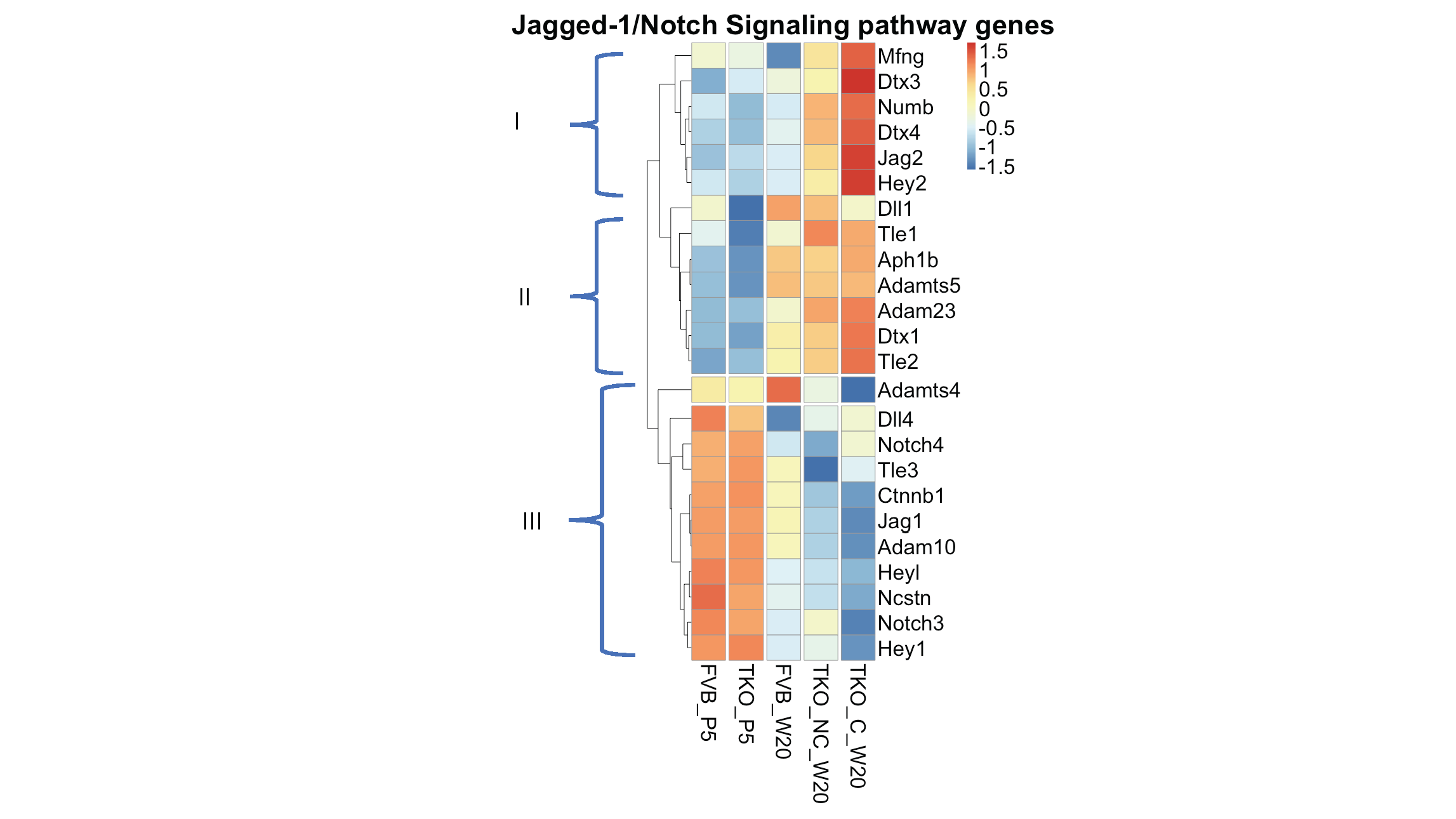

### Figure S13.tif

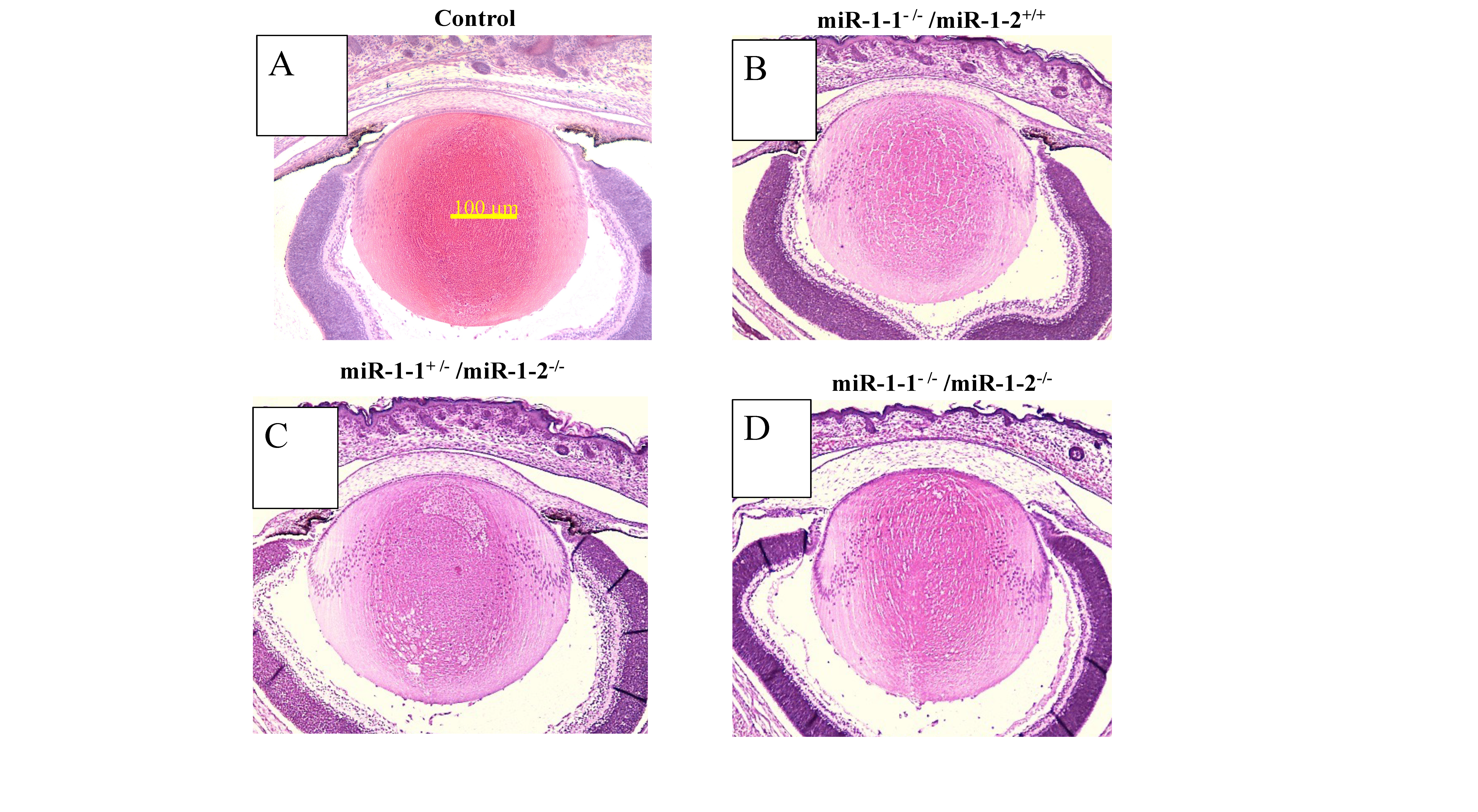

### Figure S14.tif

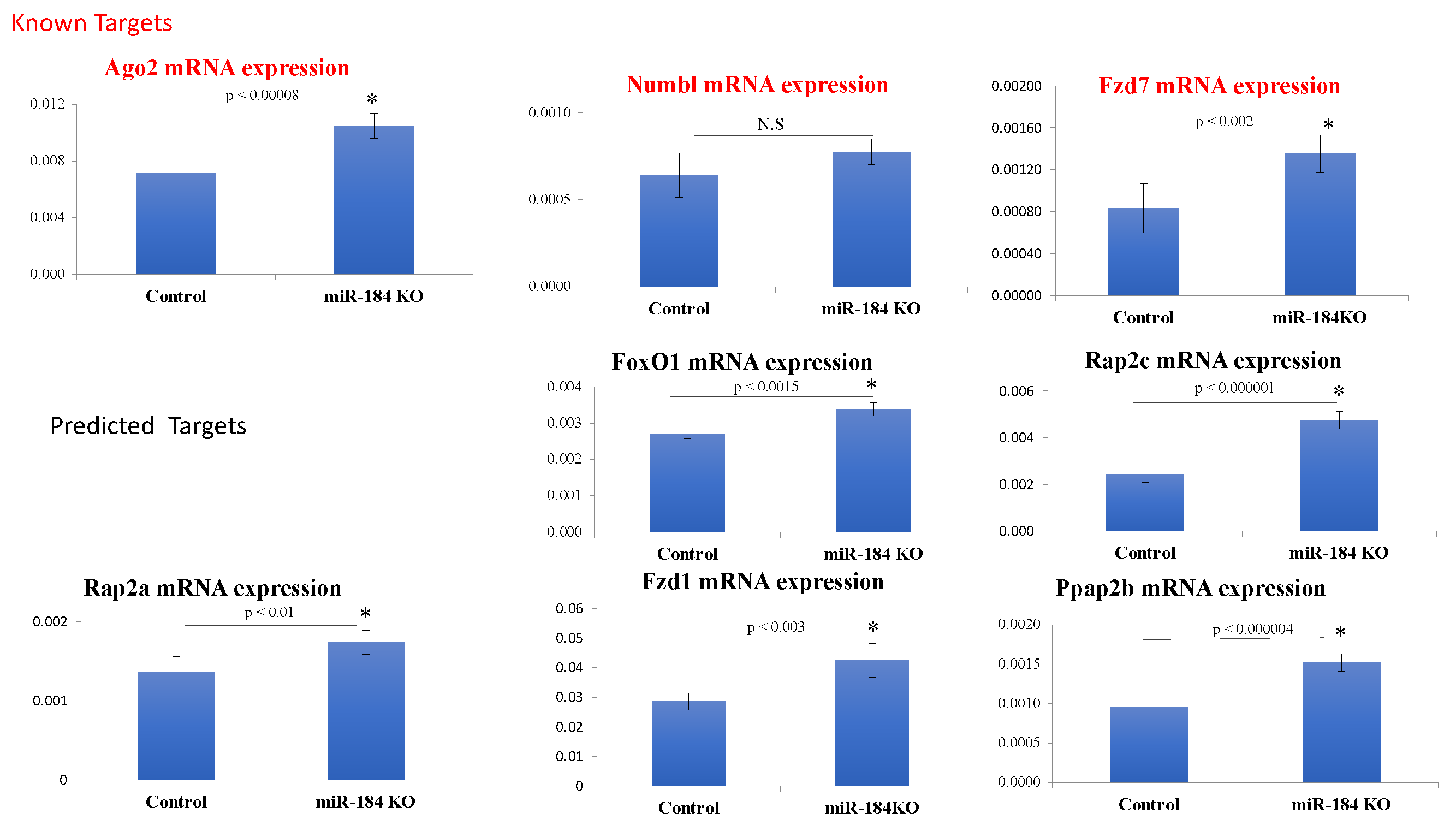
